# CD4 and CD8 co-receptors modulate functional avidity of CD1b-restricted T cells

**DOI:** 10.1101/2020.10.17.332072

**Authors:** Charlotte A. James, Yuexin Xu, Melissa S. Aguilar, Lichen Jing, Erik D. Layton, Martine Gilleron, Adriaan J. Minnaard, Thomas J. Scriba, Cheryl L. Day, Edus H. Warren, David M. Koelle, Chetan Seshadri

## Abstract

CD4 and CD8 co-receptors define distinct lineages of T cells restricted by major histocompatibility complex (MHC) Class II and I molecules, respectively. Co-receptors interact with the T cell receptor (TCR) at the surface of MHC-restricted T cells to facilitate antigen recognition, thymic selection, and functional differentiation. T cells also recognize lipid antigens presented by CD1 molecules, but the role that CD4 and CD8 play in lipid antigen recognition is unknown. We studied the effect of CD4 and CD8 on the avidity, activation, and function of T cells specific for two CD1b-presented mycobacterial lipid antigens, glucose monomycolate (GMM) and diacylated sulfoglycolipids (SGL). In a human cohort study using SGL-loaded CD1b tetramers, we discovered a hierarchy among SGL-specific T cells in which T cells expressing the CD4 or CD8 co-receptor stain with a higher tetramer mean fluorescence intensity (MFI) than CD4-CD8- T cells. To determine the role of the TCR co-receptor in lipid antigen recognition, we exogenously expressed GMM and SGL-specific TCRs in Jurkat or polyclonal T cells and quantified tetramer staining and activation thresholds. Transduced CD4+ primary T cells bound the lipid-loaded CD1b tetramer with a higher MFI than CD8+ primary T cells, and transduced CD8+ Jurkat cells bound the SGL-CD1b tetramer with higher MFI than CD4-CD8- Jurkat cells. The presence of either co-receptor also decreased the threshold for IFN-γ secretion. Further, co-receptor expression increased surface expression of CD3ε, suggesting a mechanism for increased tetramer binding and activation. Finally, we used single-cell sequencing to define the TCR repertoire and *ex vivo* functional profiles of SGL-specific T cells from individuals with M.tb disease. We found that CD8+ T cells specific for SGL express canonical markers associated with cytotoxic T lymphocytes, while CD4+ T cells could be classified as T regulatory or T follicular helper cells. Among SGL-specific T cells, only those expressing the CD4 co-receptor also expressed Ki67, suggesting that they were actively proliferating at the time of sample collection. Together, these data reveal that expression of CD4 and CD8 co-receptor modulates TCR avidity for lipid antigen, leading to functional diversity and differences in *in vivo* proliferation during M.tb disease.

## INTRODUCTION

T cells express a T cell receptor (TCR) that mediates recognition of antigens in the context of antigen presenting molecules. Most T cells recognize peptide antigens in the context of major histocompatibility (MHC) class I or class II, which express CD8 or CD4 co-receptors, respectively. TCR co-receptors facilitate positive selection and help to define functional subsets. Co-receptors also increase TCR avidity for peptide-MHC, which has downstream effects on proliferation, memory phenotype acquisition, and functional differentiation (Kersh et al., 1998; Stone et al., 2009; Viganò et al., 2012; Zehn et al., 2009). Canonically, T cells that express the CD4 co-receptor can be divided into four major functional subsets: Th1, Th2, Th17, and Treg. These functional lineages are driven by the master transcription factors T-bet, GATA3, ROR-γt, and FOXP3, respectively (Fontenot et al., 2003; Hori et al., 2003; Ivanov et al., 2006; Szabo et al., 2000; Zhen and Flavell, 1997). Conversely, peptide-specific T cells that express the CD8 co-receptor are traditionally cytotoxic T cells, which are defined by the expression of cytolytic effector molecules, such as granzymes and perforin (Apasov et al., 1993). These classifications are not absolute as T cells can exhibit functional plasticity and alter their functional program over time (Sallusto et al., 2018).

A small but consistent minority of T cells that do not express either CD4 or CD8 co-receptor provided proof-of-concept for MHC-independent modes of T cell activation (Porcelli et al., 1992). Specifically, invariant natural killer T (iNKT) cells recognize lipids presented by cluster of differentiation 1 (CD1) and mucosal associated invariant T (MAIT) cells recognize metabolites presented by MHC-related protein 1 (MR1) (Beckman et al., 1994; Kjer-Nielsen et al., 2012). On average, 15% of iNKT cells express CD4, 49% are double negative (DN), and roughly 34% express the CD8αα homodimer (O’Reilly et al., 2011). Among MAIT cells, 35% express CD8αα and 45% express CD8αβ (Gherardin et al., 2018). There is evidence that co-receptors are actively involved in antigen recognition by iNKT and MAIT cells. CD4 potentiates iNKT cell activation leading to sustained TCR signaling and potentiation of effector responses (Thedrez et al., 2007). Additionally, blocking CD8 with a monoclonal antibody leads to decreased MAIT cell responses to *Escherichia coli* (Kurioka et al., 2017). iNKT and MAIT cells can also be divided into distinct functional classes based on co-receptor expression. In humans, iNKT cells that express the CD4 co-receptor simultaneously secrete both Th1 and Th2 cytokines, and DN iNKT cells have a Th1 phenotype (Gumperz et al., 2002; Lee et al., 2002). CD8 MAIT cells express higher levels of granulysin, granzyme B, and perforin, suggesting that they are more potently cytotoxic (Dias et al., 2018). DN MAIT cells express less IFN-γ and more IL-17 than CD8 MAIT cells, and have a higher ROR-γt to T-bet ratio, indicative of a Th17 phenotype (Dias et al., 2018). Notably, these functional classes exist without two pathways for selection, antigen processing, and antigen presentation as in the case of MHC-I and MHC-II. Thus, despite early reports that suggested these “innate-like” T cells might be limited to the minority of T cells lacking expression of co-receptors, iNKT and MAIT cells are clearly part of the majority of T cells that express either CD4 or CD8.

T cells also recognize bacterial cell wall lipids presented by Group 1 CD1 molecules (CD1a, CD1b, and CD1c), and this population of T cells may be functionally distinct from iNKT and MAIT cells (Lepore et al., 2018). The contribution of TCR co-receptors in the recognition and functional differentiation of Group 1 CD1-restricted T cells is unknown. A study of T cell clones that recognize a mycobacterial cell wall glycolipid presented by CD1b, glucose monomycolate (GMM), revealed that high affinity T cell clones expressed the CD4 co-receptor and lower affinity T cell clones expressed either the CD8αβ heterodimer or were DN (Van Rhijn et al., 2014). Further, T cell clones expressing high affinity TCRs were biased towards the expression of Th1 cytokines, whereas T cell clones with lower affinity TCRs expressed Th1 and Th17 cytokines (Van Rhijn et al., 2014). These limited data suggest that TCR co-receptors may influence the recognition of lipid antigens and the functional lineage of lipid-specific T cells.

We used T cells specific for GMM and sulfoglycolipids (SGL), a class of lipids that are uniquely expressed by *Mycobacterium tuberculosis* (M.tb) and presented to T cells by CD1b, as a model system to investigate the impact of co-receptor expression on T cell function (Gilleron et al., 2004). We examine co-receptor expression on SGL-specific T cells directly *ex vivo*, as well as differences in SGL-CD1b and GMM-CD1b tetramer binding in *ex vivo* T cells, *in vitro-derived* T cell lines, and T cells transduced with an exogenous T cell receptor. We found that SGL-specific T cells identified *ex vivo* by tetramer show a bias toward CD4 co-receptor expression, and that CD4 T cells bind SGL-CD1b tetramer with a higher intensity and have lower activation thresholds than T cells that express the CD8 co-receptor or are DN. Single cell transcriptional studies of SGL-specific T cells revealed distinct TCR repertoires and functional programs that could be classified on the basis of CD4 and CD8 expression. These programs more closely align with functional subsets of peptide-specific T cells rather than iNKT or MAIT cells, revealing a previously unappreciated diversity among the functional profiles of CD1b-restricted T cells.

## RESULTS

### Ex vivo co-receptor expression by SGL-specific T cells

We first determined which co-receptor was expressed by SGL-specific T cells directly *ex vivo* using CD1b tetramers loaded with a biologically validated synthetic analog of SGL (James et al., 2018). We examined peripheral blood mononuclear cells (PBMC) derived from five U.S. healthy donors, five South African adolescents with latent tuberculosis infection (LTBI), and five South African adolescents without LTBI (Supplemental Table 1, Supplemental Table 2, and Supplemental Figure 1) (Mahomed et al., 2011). To ensure accurate identification of antigen-specific T cells, we used a dual-tetramer labelling strategy incorporating SGL-CD1b tetramer labeled with two distinct fluorochromes and defined SGL-specific T cells as staining with both tetramers while being unlabeled by a mock-loaded CD1b tetramer (Figure 1A, Supplemental Figure 1). Gates defining SGL-CD1b tetramer-positive cells were set based on staining an SGL-specific T cell line and fluorescence minus one (FMO) controls (Figure 1A) (James et al., 2018). Representative staining from one individual is shown (Figure 1B). The frequency of SGL-CD1b tetramer-positive T cells did not significantly differ between groups (p = 0.4, Supplemental Figure 1).

**Figure 1.**
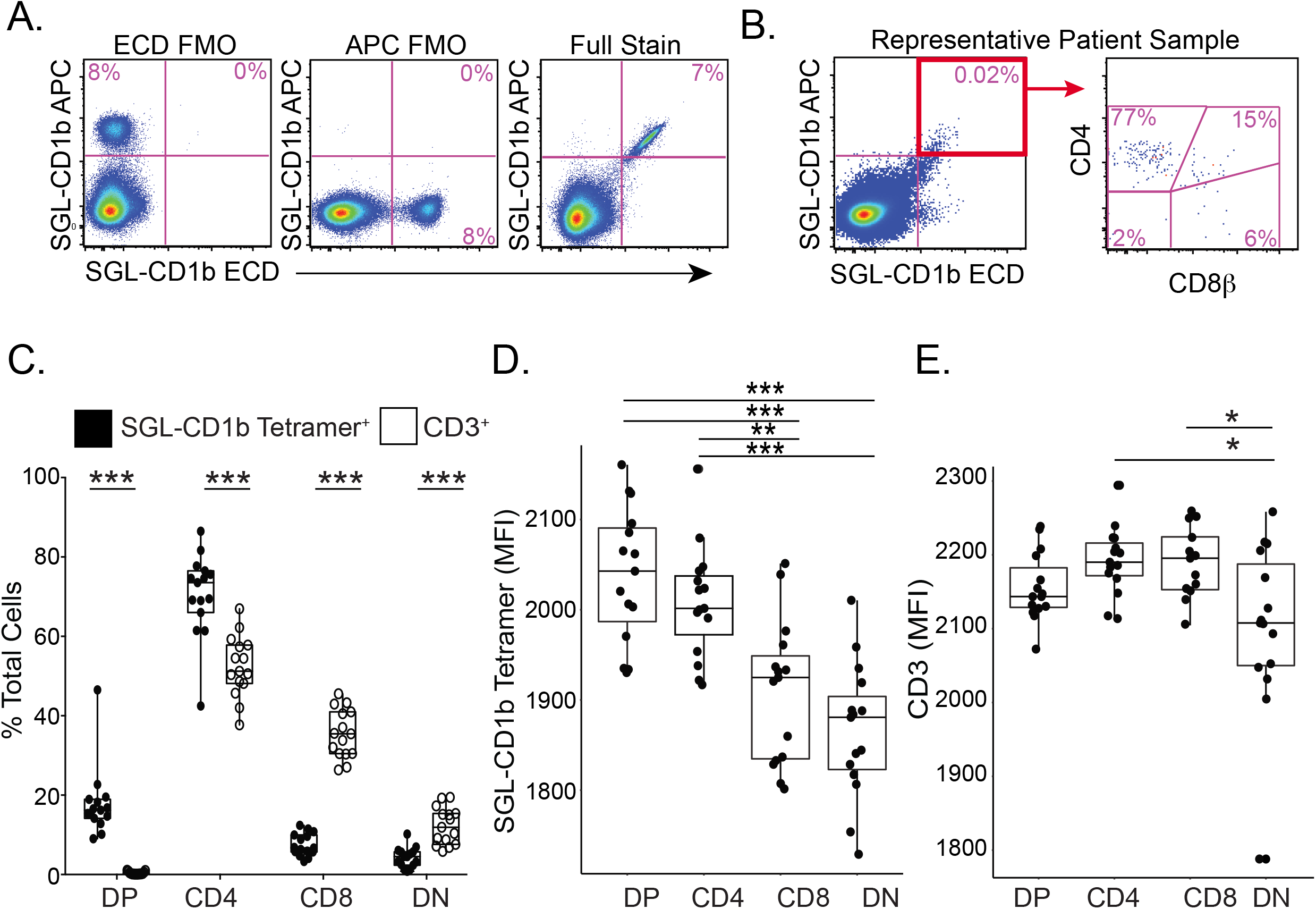
*Ex vivo* co-receptor expression by SGL-specific T cells. SGL-CD1b tetramers were incorporated into a multi-parameter flow cytometry assay to measure co-receptor expression by SGL-specific T cells. (A) The tetramer positive gate was defined by a dual tetramer staining with electron coupled dye (ECD) and allophycocyanin (APC)) and ‘Fluorescence Minus One’ (FMO) negative control (left and center) and a positive control using SGL-specific T cell line (A05) diluted in donor PBMC (right). (B) Representative staining from a South African adolescent blood donor. (C) The co-receptor expression of SGL-CD1b tetramer positive T cells in the blood was quantified using cryopreserved PBMC obtained from three groups of healthy participants: U.S. controls at low risk for M.tb exposure (n=5), South African adolescents with latent tuberculosis (IGRA-positive, n=5), and South African adolescents without latent tuberculosis (IGRA-negative, n=5). Boxplots depict the median and interquartile range of tetramer-positive cells (white) within each co-receptor group (double positive (DP), CD4, CD8, and double negative (DN), expressed as a percent of total tetramer-positive cells. Each dot represents the percent of one sample. The percent of cells in each group was compared to that present in total CD3^+^ T cells (grey). (Mann-Whitney with Bonferroni Correction, ** = p < 0.0001, n = 15). (D) Boxplots depict the median and interquartile range of mean fluorescence intensity (MFI) of SGL-CD1b tetramer-positive cells in each co-receptor group. Each dot represents the MFI of one sample. DP, CD4, and CD8 T cells were compared using Kruskal-Wallis with post-hoc Dunn test, *** = p < 0.0001, ** = 0.0001 < p < 0.001, n = 15). (E) Boxplots depict the median and interquartile range of mean fluorescence intensity (MFI) of CD3 among tetramer-positive cells in each co-receptor group. Each dot represents the MFI of one sample. MFI was compared between groups (Kruskal-Wallis with post-hoc Dunn test, * = 0.001 < p < 0.05, n = 15).

SGL-CD1b specific T cells exhibited an 18-fold enrichment of CD4 and CD8 double-positive (DP) cells (p < 0.0001) and a 1.5-fold enrichment of CD4 T cells (p < 0.0001) relative to total CD3^+^ T cells. This was accompanied by a 5.6-fold reduction of CD8 cells (p < 0.0001) and a 2.8-fold reduction of CD4 and CD8 double negative (DN) cells (p < 0.0001) (Figure 1C). Further, SGL-CD1b specific T cells expressing any combination of co-receptors exhibit consistently higher APC tetramer MFI when compared to DN SGL-CD1b specific T cells (p < 0.0001) (Figure 1D). We considered the possibility that the difference in tetramer MFI could be explained by differences in TCR expression. We found no differences in CD3ε expression among DP, CD4, and CD8 T cells (Figure 1E). However, we did detect a 4% lower CD3ε MFI among DN T cells when compared to the other groups (p = 0.02) (Figure 1E). For additional context, we quantified the level of CD3ε expression in all DP, CD4, CD8, and DN T cells from these donors and found that CD4 and DN T cells express higher levels of CD3 than CD8 T cells, which is consistent with published literature (p < 0.0001 and p = 0.004, respectively, Supplemental Figure 1) (El Hentati et al., 2010). Taken together, these data reveal a hierarchy in which SGL-specific T cells expressing the CD4 co-receptor are present in peripheral blood at the highest frequency and stain with SGL-CD1b tetramer at the highest MFI, followed by CD8 and DN T cells, respectively.

### Co-receptors enhance functional avidity of an SGL-specific TCR

Several factors affect TCR avidity for antigen, including affinity for the antigen presenting molecule, the level of TCR expression, and the expression of CD4 and CD8 co-receptors on the cell surface (Laugel et al., 2007; Viganò et al., 2012; Viola and Lanzavecchia, 1996). The aggregate impact these factors have on the binding strength of the TCR in live cells is known as ‘functional avidity’ due to the multivalent and multifaceted nature of this interaction (Viganò et al., 2012). Our *ex vivo* data suggests SGL-specific T cells expressing the CD4 co-receptor may have higher functional avidity than those expressing CD8. To test this hypothesis directly, we examined SGL-specific T cell lines derived from South African adults with LTBI (James et al., 2018). Virtually all of the SGL-CD1b tetramer staining cells within the A01 T cell line express a heterodimeric CD8αβ co-receptor in contrast to the A05 T cell line, which was strongly biased toward CD4 expression (Supplemental Figure 2). A05 exhibited 5.5-fold higher SGL-CD1b tetramer MFI compared to A01 (p = 0.03) (Supplemental Figure 2).

We considered the possibility that the observed differences in tetramer MFI between A01 and A05 might be due to differences in their TCRs. Indeed, A01 and A05 each express a distinct TCR, which may have different affinities for the CD1b-SGL complex independent of co-receptor (Table 1). To address this, we cloned and transduced polyclonal T cells isolated from a blood bank donor with an A05 TCR expression construct (Figure 2A). The constant region of the TCR in the expression construct was replaced with a murine TCR constant region to allow quantification of exogenous TCR expression. Among successfully transduced T cells, as demonstrated by expression of the exogenous TCR and defined by staining with anti-murine TCR-β chain constant region (mTCRBC), 88% also co-stained with SGL-CD1b tetramer, revealing that TCR expression is sufficient to confer SGL-CD1b tetramer binding (Figure 2A). To further demonstrate the specificity of staining using SGL-CD1b tetramer we also transduced a TCR that is specific for the mycobacterial glycolipid glucose monomycolate (GMM) into Jurkat cells (Van Rhijn et al., 2013) (Supplemental Figure 3). These GMM-specific Jurkat cells avidly bind GMM-CD1b tetramer, whereas they largely fail to bind SGL-CD1b tetramer and mock-loaded CD1b tetramer (Supplemental Figure 3). We attempted to also express the A01 TCR in primary T cells, but exceptionally high levels of TCR expression were required for successful tetramer staining (data not shown). These data support our hypothesis that the difference in activation thresholds between A01 and A05 may be largely due to their TCRs. However, the use of transduced polyclonal T cells containing CD4, CD8, and DN populations allowed us to specifically study the role of co-receptor independent of the TCR.

**Figure 2.**
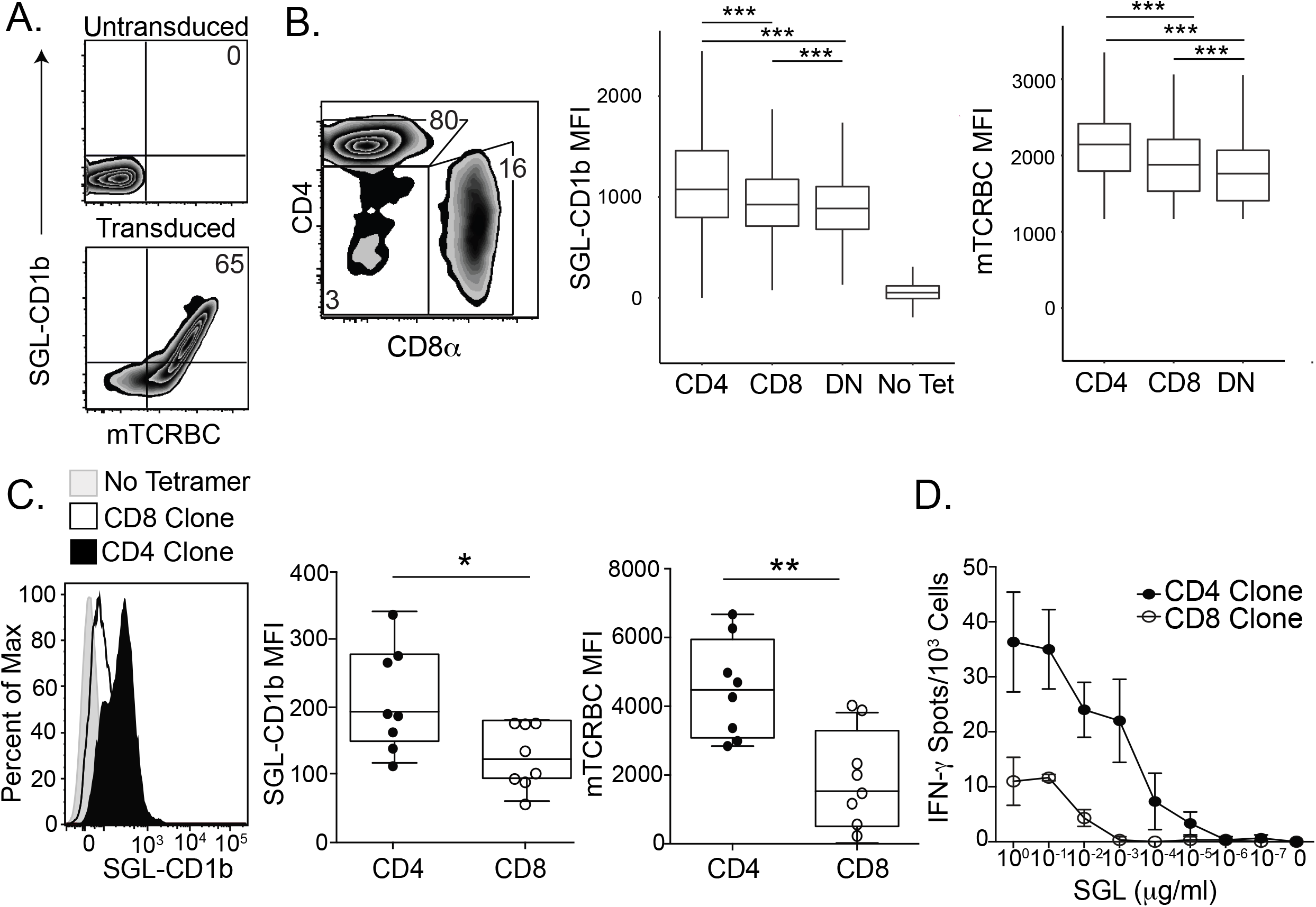
Co-receptors enhance functional avidity of an SGL-specific TCR. (A) Primary T cells were transduced with the TCR from the A05 T cell line using a lentiviral vector. Among primary T cells that are not transduced with the exogenous TCR, 0% stain with SGL-CD1b tetramers or a mTCRBC antibody (top). After transduction with the TCR, the cells stain with both the SGL-CD1b tetramer and mTCRBC antibody (bottom). (B) Flow plot depicts the percent of transduced T cells that are CD4, CD8, or DN. Boxplot depicts the median and interquartile range of the SGL-CD1b or mTCRBC MFI of each CD4, CD8, and DN T cell that was transduced with the A05 TCR (Two-way ANOVA, post-hoc Dunn test, *** = p < 0.0001, n = 86,520). Data are representative of two independent rounds of primary T cell transduction with the TCR of the A05 T cell line. (C) Single cell clones were generated from the primary T cell transductants by limiting dilution cloning. A representative CD4 (black) and CD8 (white) T cell clone stained with SGL-CD1b tetramer is compared (left). A no tetramer control is also represented (grey) (left). Boxplots depict the median and interquartile range of SGL-CD1b MFI of CD4 (black) and CD8 (white) clones (Mann-Whitney test, p = 0.019, n = 16). (D) Antigen-specific activation of CD4 (black) and CD8 (white) T cell clones was measured using IFN-γ ELISPOT. Error bars represent standard deviation of triplicate wells. SGL was serially diluted log-fold from 1 μg/ml to 10^-7^ μg/ml. Data are representative of two independent experiments.

**Table 1.** TCRs expressed by SGL-specific T Cell Lines. Clonality was determined by via staining with antibodies targeting human TCR-β variable genes (IOTest BetaMark Kit, Beckman Coulter) or high-throughput sequencing (ImmunoSEQ, Adaptive Biotechnologies). The full-length TCR sequence was determined by template-switched PCR after sorting tetramer-positive T cells. The T cell receptor gene usage and CDR3 region amino acid sequence is summarized here. Variable and joining gene segment names were assigned using the International ImMunoGeneTics (IMGT) Database.

| Lines | TCR | Variable | CDR3 | Joining |
| --- | --- | --- | --- | --- |
| **A01** | $\alpha$ | TRAV8-6 | CAVKAGYSSASKIIF | TRAJ3 |
| | $\beta$ | TRBV12-4 | CASKGKQGPEQFF | TRBJ2-1 |
| **A05** | $\alpha$ | TRAV13-2 | CAEVLGGTSYGKLTF | TRAJ52 |
| | $\beta$ | TRBV20-1 | CSASGRRGYGYTF | TRBJ1-2 |

Among T cells transduced with the A05 TCR, those also expressing CD4 stained with the highest tetramer MFI, followed by CD8 and DN T cells (p < 0.0001) (Figure 2B). This relative hierarchy was also noted within mTCRBC suggesting that the level of exogenous TCR expression post-transduction also varies based on co-receptor expression (p < 0.0001) (Figure 2B). To explore whether these findings were generalizable, we repeated these experiments using a CD1b-restricted TCR specific for glucose monomycolate (GMM) (Van Rhijn et al., 2013). We again observed a hierarchy in which transduced CD4+ T cells stained with the highest MFI and expressed the highest levels CD3ϵ followed by CD8 and DN T cells (Supplemental Figure 4). We subjected the T cells transduced with the A05 TCR to limiting dilution cloning in an attempt to generate a panel of clones with equivalent levels of exogenous TCR expression. However, we noted that the CD4 clones consistently stained at a higher MFI with SGL-CD1b tetramer than CD8 clones (p = 0.019) and also expressed higher levels of mTCRBC (p = 0.005) (Figure 2C). These data suggest that CD4 T cells may intrinsically express higher levels of exogenous TCR than CD8 T cells or possess an intrinsic advantage in their ability to be transduced with lentiviral vectors. Finally, we assessed differences in T cell activation between CD4 and CD8 T cells transduced with the A05 TCR by measuring IFN-γ production in the presence of titrating amounts of SGL. An A05-CD4 T cell clone displayed an EC50 of 0.0006 μg/ml, which was 100-fold lower than an A05-CD8 T cell clone (Figure 2D). Notably, the clones used in this assay also exhibit differences in SGL-CD1b tetramer bindings and TCR expression (Figure 2C). Taken together, these data demonstrate that an increase in tetramer binding and TCR expression confers enhanced sensitivity to activation at limiting antigen concentrations than CD8 T cells, despite expressing the same T cell receptor.

### CD8 is sufficient to enhance functional avidity of an SGL-specific TCR

Primary T cells expressing CD4, CD8, or neither co-receptor exhibit other biological differences besides their co-receptor that might affect functional avidity for SGL-CD1b complex (Zapata et al., 2004). To address this, we transduced the A05 TCR into a DN Jurkat T cell line and an isogenic T cell line that had been stably transduced to express CD8αα. As with primary T cells, the A05 TCR was sufficient to confer SGL-CD1b tetramer staining in Jurkat T cells (Figure 3A). The SGL-CD1b tetramer MFI was 1.59-fold higher among CD8αα expressing Jurkat T cells compared to DN T cells (p = 0.03) (Figure 3B). Notably, mTCRBC expression was also 1.74-fold higher among CD8αα compared to DN T cells, indicating higher levels of A05 TCR expression (p = 0.03) (Figure 3B). To investigate this further, we analyzed SGL-CD1b MFI after stratifying CD8αα expression into logarithmic tertiles (Figure 3C, left). The CD8^hl^ Jurkat cells had the highest MFI, which was 1.32-fold higher than CD8^med^ Jurkats and 2.35-fold higher than CD8^lo^ Jurkats (p = 0.0002) (Figure 3C, middle). When we compared the level of mTCRBC expression, we found that the CD8^hi^ Jurkat cells again had the highest MFI, followed by the CD8^med^ and CD8^lo^ Jurkats (p = 0.0002) (Figure 3C, right). These data support our hypothesis that co-receptor expression is sufficient to increase functional avidity for SGL-CD1b tetramer, and one mechanism by which this occurs is by increasing the expression of the TCR complex at the cell surface.

**Figure 3.**
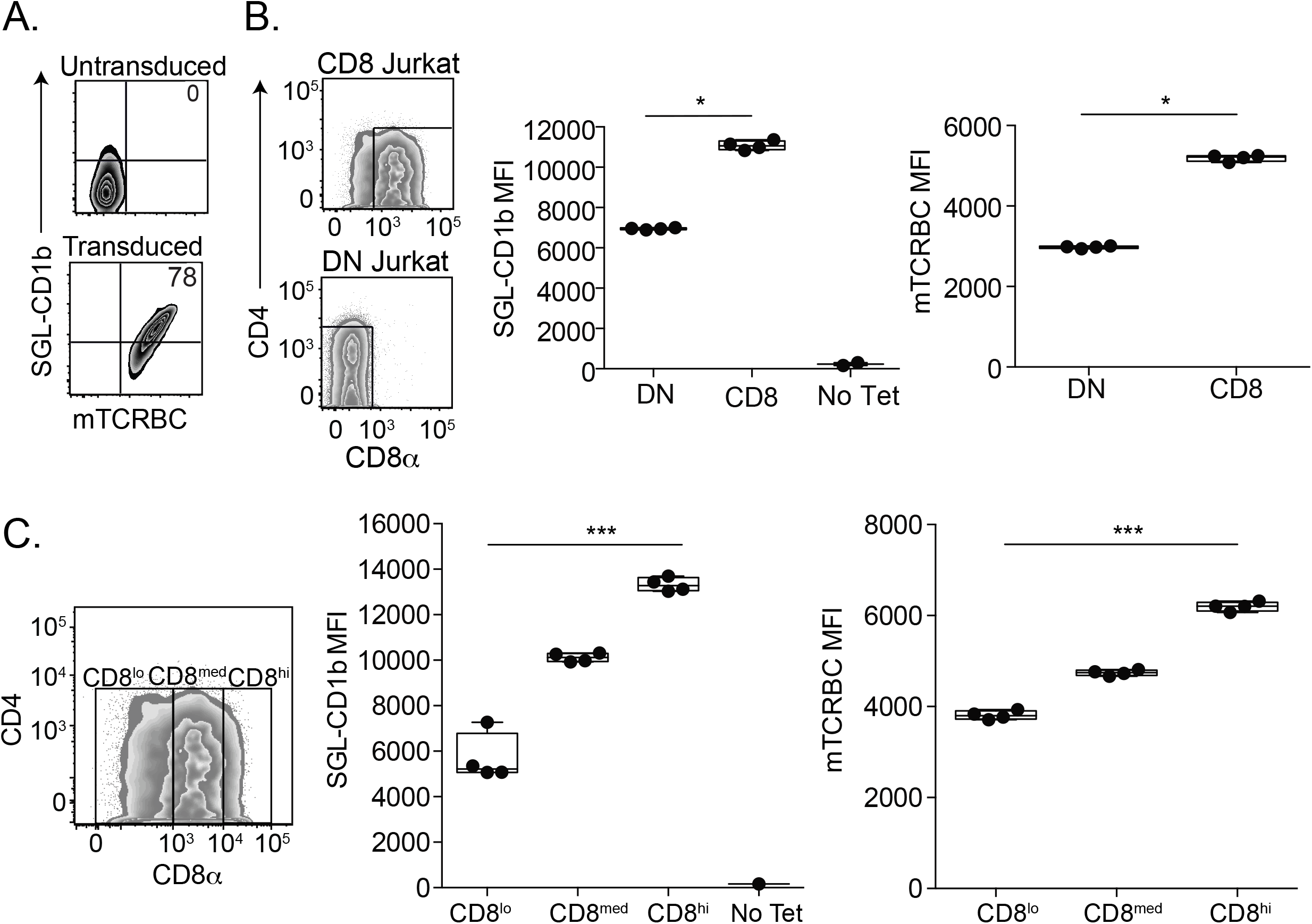
CD8 is sufficient to enhance functional avidity of an SGL-specific TCR. (A) DN and CD8 Jurkat cell lines were transduced with the TCR from the A05 T cell line. Untransduced CD8 Jurkat cells do not stain with SGL-CD1b tetramer or a murine T cell receptor chain constant region (mTCRBC) specific antibody (top). After transduction with the TCR, the cells stain with both the SGL-CD1b tetramer and mTCRBC antibody (bottom). (B) CD8 and DN Jurkat cells express a gradient of TCR co-receptor expression. Gates enclose true “CD8” (left, top) and “DN” (left, bottom) cells that express only CD8 or no co-receptors, respectively. Of note, the Jurkat cells that express CD8 display a gradient of expression, and the DN Jurkat cells exhibit low levels of CD4 co-receptor expression. The distributions of SGL-CD1b MFI of the DN (dark grey) and CD8 (white) Jurkat cells were then compared (middle) (Student’s t-test, p < 0.0001, n = 29,195). A no tetramer control is also represented (light grey) (right). The distributions of mTCRBC MFI of the DN (dark grey) and CD8 (white) Jurkat cells were also compared (right) (Student’s t-test, p < 0.0001, n = 29,195). (C) The distributions of SGL-CD1b MFI of CD8^hi^ (dark grey, CD8 MFI of 10^4^ to 10^5^), CD8^med^ (grey, MFI of 10^3^ to 10^4^) and CD8^lo^ (light grey, MFI of 0 to 10^3^) Jurkat cells were then compared (right) (Student’s t-test, p < 0.0001, n = 20,260). A no tetramer control is also represented (grey) (right). The distributions of mTCRBC MFI of CD8^hI^, CD8^med^, and CD8^lo^ Jurkat cells were then compared (right) (Student’s t-test, p < 0.0001, n = 20,260). Data are representative of three independent experiments.

### Glycolipid-specific T cells express a diverse TCR repertoire

To profile CD4 and CD8 glycolipid-specific T cells directly *ex vivo*, we sorted single cells from four South African adults with newly diagnosed active TB disease using SGL-CD1b and GMM-CD1b tetramers (Day et al., 2011). In this experiment, we included GMM-specific T cells and T cell lines with known TCRs as controls to verify our tetramer-sorting strategy. Gates defining CD1b tetramer-positive cells were set based on control samples not stained with the tetramers as well as SGL- and GMM-specific T cell lines (Supplemental Figure 1). To validate our sorting strategy, we first assessed the TCRs recovered from the T cell lines with a known TCR. We note that the V genes, J genes, and CDR3s recovered from these wells match the known TCRs (Supplemental Data 1). In addition, as several studies highlighted the enrichment of TRAV1-2 among GMM-specific TCRs we next compared the frequency of TRAV1-2 among GMM-CD1b tetramer sorted cells and bulk T cells from that same individual to verify the tetramer sorting strategy reliably isolated lipid antigen-specific T cells (DeWitt et al., 2018; Lopez et al., 2020; Van Rhijn et al., 2013). We found that 10% of recovered TCRs express TRAV1-2, compared to 2.9% of bulk T cells (p = 0.0005) (Supplemental Figure 5, Supplemental Table 3). We were also able to identify TCR V genes that have been described within diverse GMM-specific TCR repertoires, such as TRAV17, TRAV8, TRAV12, and TRAV13 (Supplemental Figure 5) (DeWitt et al., 2018; Lopez et al., 2020; Van Rhijn et al., 2013).

As relatively little is known about SGL-specific TCRs, we first compared the recovered TCR variable genes from cells sorted using the SGL-CD1b tetramer to the TCRs from our A01 and A05 T cell lines (Table 1). 4.7% of recovered TCRs from SGL-CD1b tetramer-sorted cells used TRAV8-6 (A01) and 1.6% used TRAV13-2 (A05), compared to 2.1% and 1.0% of bulk T cells, respectively (p = 0.053 and p = 0.35, respectively) (Supplemental Figure 5, Supplemental Table 3). We also found that approximately 6.2% of TCRs recovered from SGL-CD1b tetramer-sorted cells used TRAV21 and 6.2% used TRAV19, compared to 2.9% and 4.2% of bulk T cells, respectively (p = 0.031 and p = 0.26, respectively) (Supplemental Figure 5, Supplemental Table 3). These data reveal that SGL-specific T cells isolated and examined directly *ex vivo* express diverse TCRs that likely have varying functional avidities for CD1b.

Next, we examined the TCR-a chain gene usage of CD4 and CD8 SGL-specific T cells to determine whether the TCR repertoires were distinct between these subpopulations (n = 38 and n = 43, respectively). Overall, 30% of the V genes we detected were found in both CD4 and CD8 SGL-specific T cells (Supplemental Figure 6). Of the genes that were found in more than one TCR, TRAV21 and TRAV12-2 were detected preferentially among CD8 SGL-specific T cells (p = 0.06 and p = 0.12, respectively) (Supplemental Figure 6). We also found that TRAV2, TRAV29, and TRAV9-2 were detected preferentially among CD4 SGL-specific T cells (p = 0.10, p = 0.21, p = 0.21, respectively) (Supplemental Figure 6). These data suggest that the TCR repertoire of CD4 and CD8 SGL-specific T cells may be distinct.

### CD4 and CD8 identify functionally distinct compartments among glycolipid-specific T cells

To examine whether the expression of CD4 or CD8 co-receptors was associated with different functional profiles, we used single cell sequencing to examine the transcriptional profiles of GMM- and SGL-specific T cells (n = 223), tetramer-negative T cells (n = 43), and T cell lines as positive controls (n = 6) (Han et al., 2014). In an unsupervised analysis using a distance-based clustering approach, we found seven clusters that were represented equally among both tetramer-positive and tetramer-negative T cells (Supplemental Figure 7) (Satija et al., 2015). We then performed a supervised analysis focusing on cells in which either CD4 or CD8 transcript was detected (105 of 272 total cells) (Supplemental Figure 7). We identified three distinct clusters using Uniform Manifold Approximation and Projection (UMAP) (Figure 4A) (McInnes et al., 2018). T cells expressing the CD4 and CD8 co-receptor were spatially separated, suggesting that they express unique transcriptional profiles (Figure 4B).

**Figure 4.**
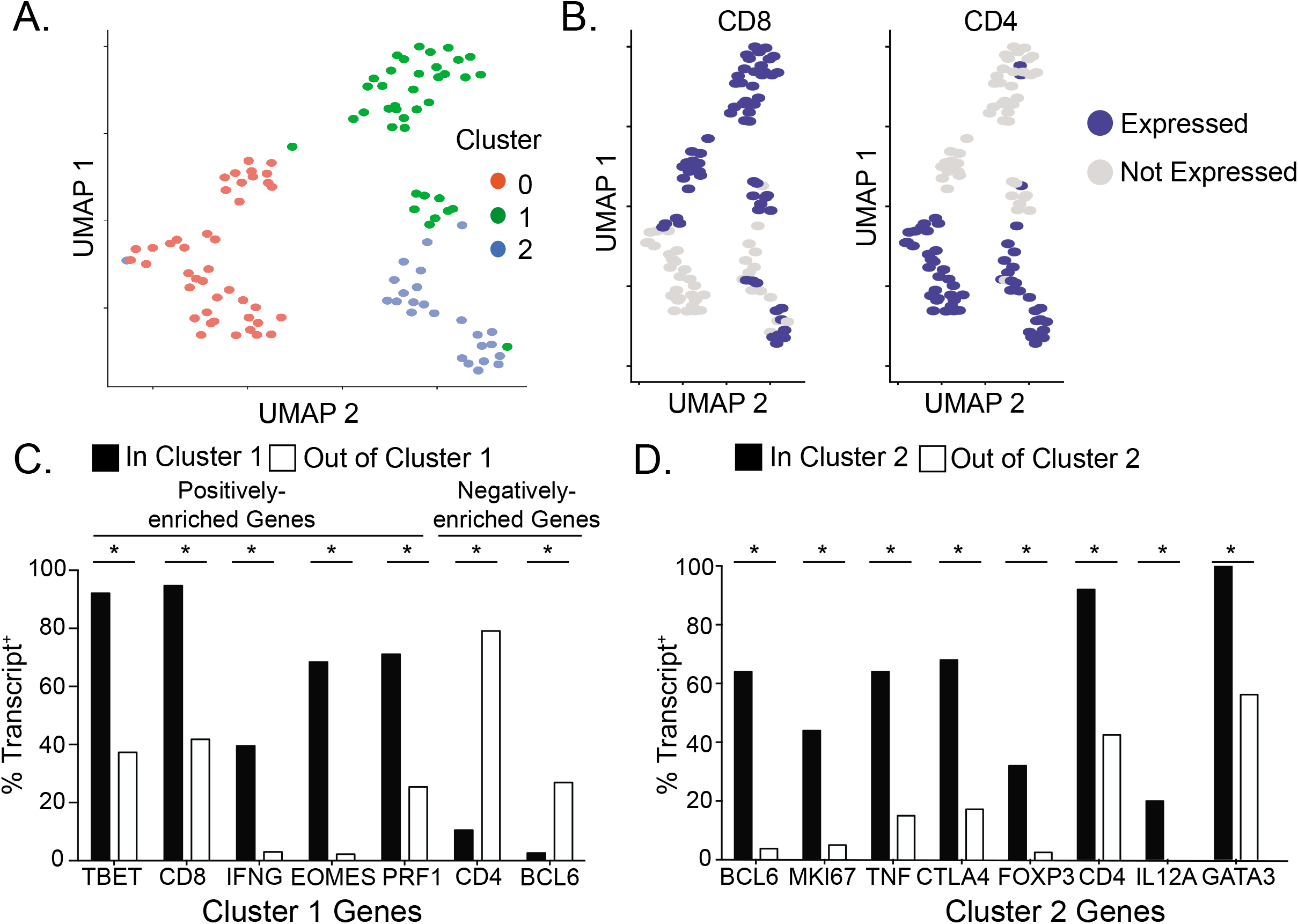
CD4 and CD8 identify functionally distinct compartments among glycolipid-specific T cells. SGL-CD1b and GMM-CD1b tetramers were incorporated into a multi-parameter flow cytometry assay to isolate SGL-specific and GMM-specific T cells using fluorescence activated cell sorting (FACS). (A) Functional profiles of sorted cells were visualized in two-dimensional space using the dimensionality reduction method UMAP. Each dot represents one cell. The three distinct clusters are represented in red, blue, and green (n = 105). (B) UMAP visualization of sorted cells colored by CD8 (left) and CD4 (right) expression. Cells that express the respective co-receptor are colored blue and cells that do not express the co-receptor are colored grey. Bars represent the percent of cells within (black) and outside (white) of (C) cluster 1 or (D) cluster 2 that express the listed genes. Each listed gene is statistically positively or negatively enriched within cluster 1 (p-values listed in Supplemental Table 4, Wilcoxon Rank Sum Test with Bonferroni correction, n = 105).

To determine which genes were driving the differences between the clusters, we used an established approach for differential gene expression analysis in single-cell RNA sequencing experiments (Satija et al., 2015). Cluster 1 is enriched for CD8 T cells that express Th1 and cytotoxic transcripts such as, TBET, IFN-γ, EOMES, and PRF1 (Figure 4C, Supplemental Table 4). Cluster 2 is characterized by a strong enrichment of CD4 T cells that express a variety of functional transcripts, including Th1 (TNF and IL-12A), Th2 (GATA3), regulatory (CTLA4 and FOXP3), and the TFH transcription factor, BCL6 (Figure 4D, Supplemental Table 4). No genes were positively enriched in Cluster 0, and genes positively enriched in cluster 1 and cluster 2 were negatively enriched in cluster 0 (PRF1, TBET, EOMES, CD8, GATA3, IFNG, BCL6) (Supplemental Table 4).

These data were complemented by comparing the functional profiles of our *in vitro*-derived T cell lines, A01 and A05, which express the CD8 and CD4 co-receptor, respectively (Supplemental Figure 2). We profiled cytokine expression using a previously optimized intracellular cytokine staining panel in the presence of SGL and K562 antigen presenting cells that have been stably transfected to express CD1b (K562-CD1b) or empty vector (K562-EV) (de Jong et al., 2010; De Rosa et al., 2012). The majority of cells from both T cell lines expressed IFN-γ, TNF, and IL-2 in the presence of SGL (Figure 5A). In contrast, the T cell lines diverged in the expression of CD40L and CD107a, which are markers of B cell help and degranulation, respectively (p = 0.03 and p = 0.03, respectively) (Figure 5A). A01 also secreted 10-fold more granzyme B than A05 and specifically lysed 67% of target cells in the presence of antigen, compared to −2% lysis when no antigen was present (p = 0.03) (Figures 5B, 5C, and 5D). By comparison, A05 did not exhibit cytolytic activity in the presence of antigen above the condition with no antigen present (p = 0.4) (Figure 5C and 5D). Taken together, these data support the conclusions of our single cell transcriptional profiling experiments and show that SGL-specific T cells expressing CD4 generally express cytokines that align with a T-helper phenotype, while those expressing CD8 have a cytotoxic effector phenotype.

**Figure 5.**
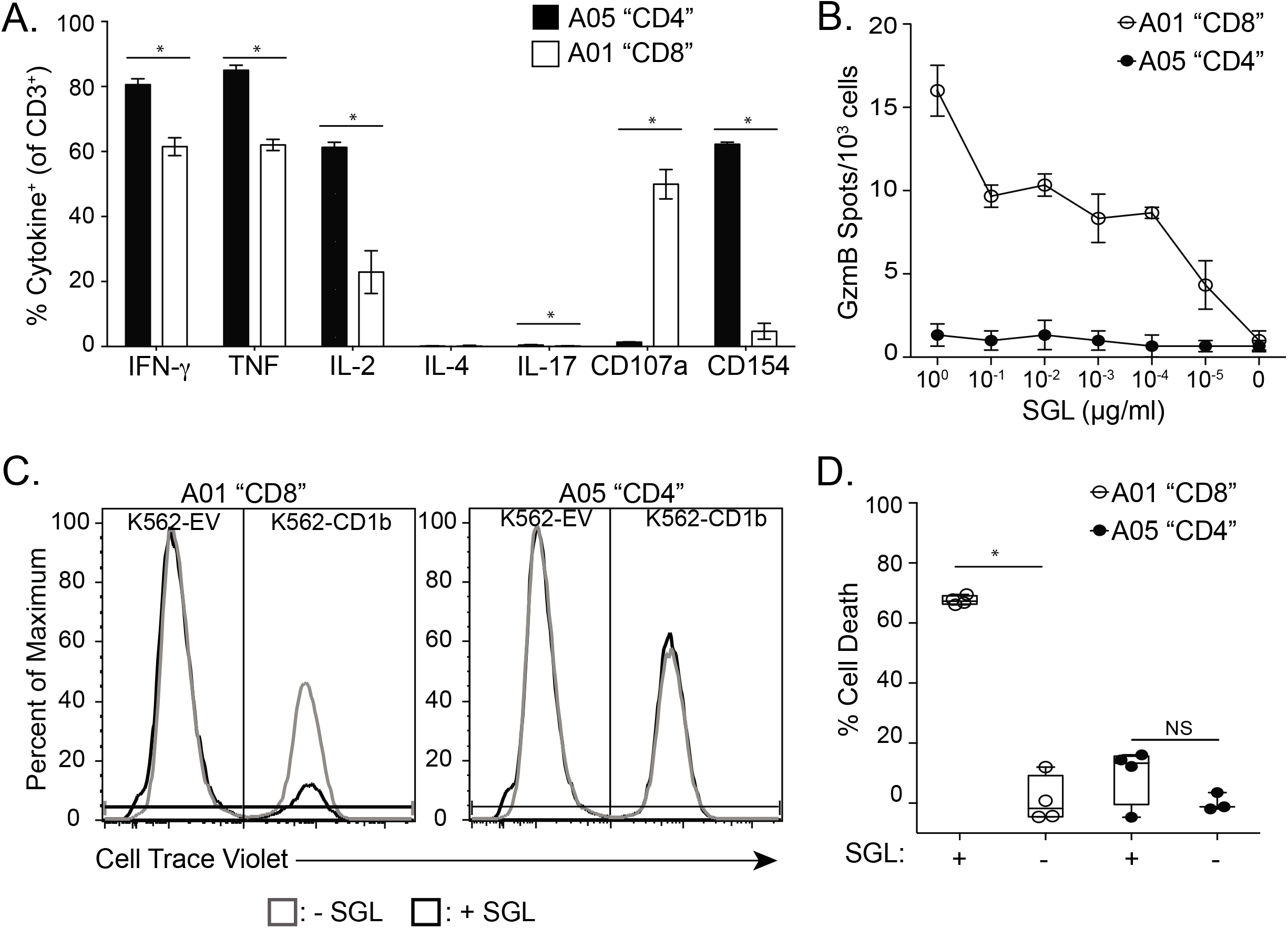
Functional profiles of the A01 and A05 T cell lines. (A) Cytokine production by the A01 (white) and A05 (black) T cell lines was measured using intracellular cytokine staining after co-incubation with K562-CD1b or K562-EV antigen presenting cells and SGL antigen for 6 hours. Data are expressed as a percent of T cells that express the cytokine as a percentage of total CD3+ T cells. Percentages are background corrected by subtracting the percentage of T cells that express cytokine after co-incubation with K562-EV antigen presenting cells and SGL for 6 hours. Error bars represent the mean and standard deviation of four technical replicates. (* = p = 0.03, Mann Whitney, n = 8). Data are representative of at least two independent experiments. (B) Granzyme B (GzmB) production from 1,000 T cells from the A01 (white) and A05 (black) T cell lines after co-incubation with K562-CD1b antigen presenting cells and SGL antigen. Cells were cultured overnight and GzmB production was assessed using GzmB ELISPOT. Antigen concentration ranged from 1 μg/ml to 10^-5^ μg/ml in log-fold dilutions. Error bars represent standard deviation of triplicate wells. Data are representative of two independent experiments. (C) Cytotoxicity was assayed by co-incubating A01 (left) and A05 (right) T cell lines with K562-EV and K562-CD1b antigen presenting cells labeled “low” and “high” with Cell Trace Violet, respectively. Co-cultures were incubated in the presence (red) or absence (blue) of SGL antigen. We defined specific lysis as the ability of a T cell line to specifically reduce the K562-CD1b cell population in the presence of lipid, while leaving the K562-EV population and the K562-CD1b without pulsed antigen intact after co-incubation. (D) Cytotoxicity was quantified by calculating the percent of cell number reduction of K562-CD1b cells in the experimental conditions compared to the percentage of K562-CD1b cells in culture in a condition with no T cells and no lipid antigen (Supplemental Figure 8) (% Cell Death). Percentages were calculated for the A01 (white) and A05 (black) T cell lines in the presence or absence of SGL. Error bars represent the mean and standard deviation of four technical replicates (* = p = 0.03, Mann Whitney, n = 8). Data are representative of at least two independent experiments.

Finally, we examined co-expression of specific transcripts from SGL- and GMM-specific T cells. Among both tetramer-positive and tetramer-negative T cells, we identified T cells that co-expressed the transcription factors TBET and GATA3, suggestive of functional plasticity (r = 0.29, p = 0.003) (Fang and Zhu, 2017) (Figure 6A). The transcription factors RORC and RUNX3, which are canonically expressed by Th17 and cytotoxic peptide-specific T cells, respectively, were ubiquitously expressed (Figure 6A) (Ivanov et al., 2006; Shan et al., 2017). There was also a significant correlation between CTLA4 and FOXP3, indicating that these two markers are co-expressed by CD4 glycolipid-specific T cells that may have a regulatory function (r = 0.36, p = 0.0001) (Figure 6B). MKI67, which is expressed by actively proliferating cells, was enriched in the same cluster as CD4 (Figure 4D, Figure 6C, Supplemental Table 4) (Scholzen and Gerdes, 2000) with 44% of T cells in Cluster 2 expressing MKI67, whereas only 5% of T cells in Cluster 0 and Cluster 1 express this marker (Figure 4D, Figure 6C) (p < 0.0001, Supplemental Table 4). These data suggest that SGL-specific CD4 T cells, but not CD8 T cells, may have been actively proliferating at the time of sample collection, perhaps as a result of increased functional avidity in the setting of limited antigen availability. Of note, MKI67 was expressed in 15.1% of CD1b-restricted T cells that were included in our final analysis and 0% off tetramer-negative T cells, which suggests that Ki-67 expression is not generally expressed in peripheral blood T cells in individuals with active TB (p = 0.12).

**Figure 6.**
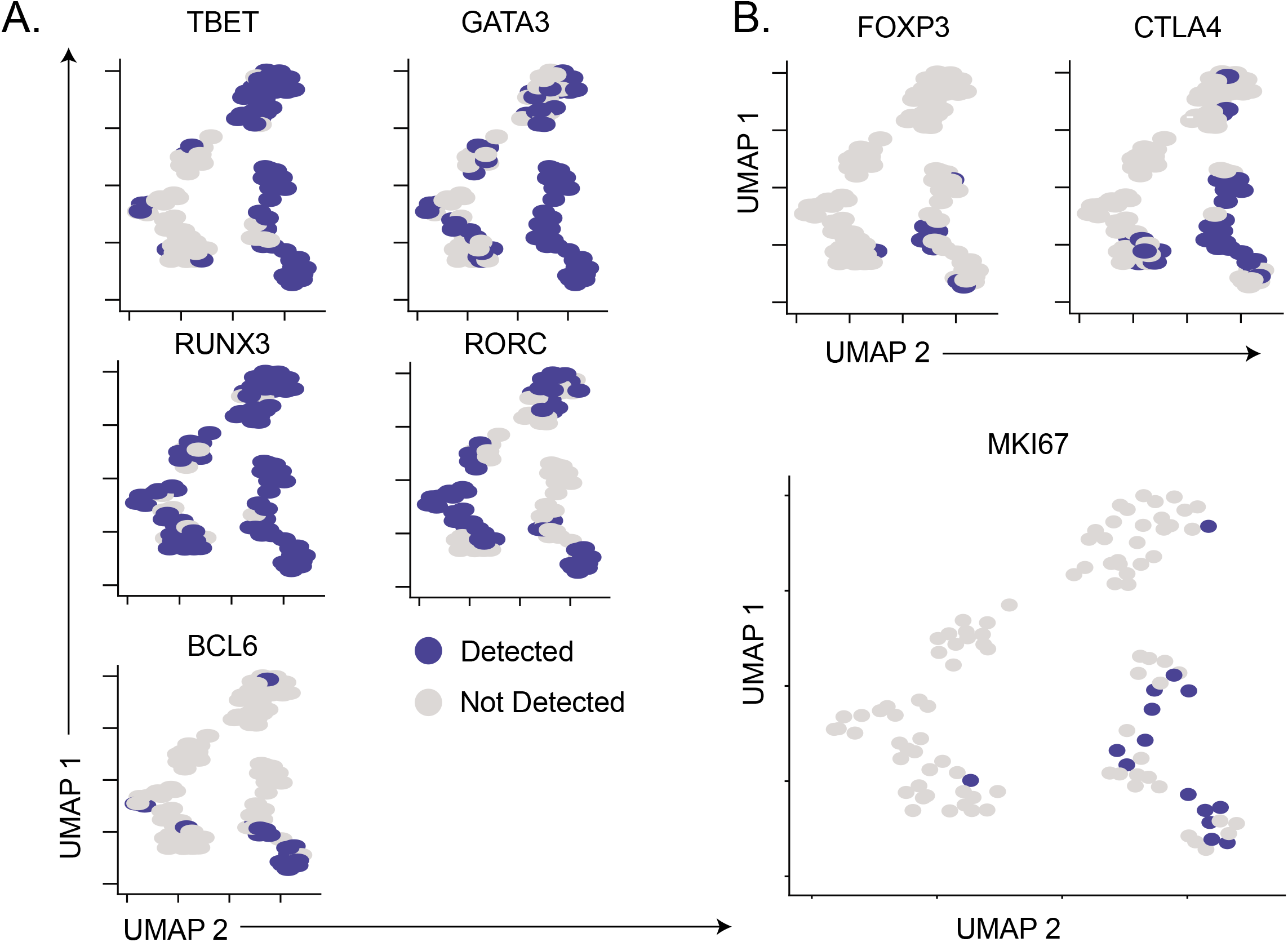
Co-expression of transcription factors and cellular markers among glycolipid-specific T cells. (A) Functional profiles of SGL-CD1b- and GMM-CD1b-specific T cells isolated from two individuals with active TB cells were visualized in two-dimensional space using the dimensionality reduction method UMAP. Each dot represents one cell. UMAP visualization of sorted cells is colored by TBET (top left), GATA3 (top right), RORC (middle left), BCL6 (middle right), FOXP3 (bottom left), and CTLA4 (bottom right). Cells that express the respective transcript are colored blue and cells that do not express the co-receptor are colored grey (n = 105). (B) UMAP visualization of sorted cells is colored by MKI67 expression. Cells that express the respective transcript are colored blue and cells that do not express the co-receptor are colored grey (n = 105).

## DISCUSSION

In summary, we have shown that SGL-specific T cells analyzed directly *ex vivo* exhibit a bias towards CD4 expression, and our data reveal that CD4 SGL-specific T cells have a higher functional avidity than CD8 SGL-specific T cells. Whether we examined *ex vivo, in vitro-derived*, or transduced SGL-specific T cells, we observed a consistent hierarchy in which CD4 T cells bind SGL-CD1b tetramer with a higher fluorescence intensity and have the lowest activation thresholds, followed by CD8 T cells, and DN T cells. In fact, our data show that the surface expression of a TCR co-receptor is associated with increased levels of TCR expression, revealing a potential mechanism for the increase in activation and functional tetramer avidity we observed. Finally, we examined the ex vivo transcriptional profiles of SGL-specific T cells in patients with active TB and find canonical transcription factor lineages among CD4 and CD8 CD1b-restricted T cells. These data significantly advance our understanding of how coreceptors modulate the recognition of lipid antigens and the functional profiles of CD1b-restricted T cells in humans.

Our data contrast with a previous report that the CD4 co-receptor is not involved in lipid antigen recognition by CD1b-restricted T cells (Sieling et al., 2000). This study used blocking antibodies targeting the interaction between CD4 and MHC-II, which may not have reliably predicted the effect on CD1b. Further, the use of co-receptor blocking antibodies can have pleotropic effects, including paradoxical activation, of T cells (Campanelli et al., 2002). We adopted a molecular approach to this question, and our results are consistent with previous reports that the CD4 co-receptor enhances iNKT cell activation (Chen et al., 2007; Thedrez et al., 2007). Our data extend these studies by revealing a potential molecular mechanism in the form of stabilized TCR expression at the surface of CD1b-restricted T cells, which would not have emerged from experiments using blocking antibodies.

In addition to increasing TCR expression, there may be other potential mechanisms that contribute to increasing the functional avidity of CD1b-restricted T cells, and these mechanisms are not mutually exclusive. First, TCR co-receptors may physically bind to CD1b and augment TCR affinity (Janeway, 1992). However, due to the structural differences between MHC-II and CD1b, putative binding sites for this interaction are not obvious. Second, precise spatial control of the immunologic synapse is important for tuning TCR sensitivity to antigen, possibly by controlling the level of cholesterol in lipid rafts (Kabouridis, 2006; Roh et al., 2015; Schieffer et al., 2014). Third, the rate of recycling in endosomes and activation state of T cells can influence T cell receptor stability at the cell surface (Favier et al., 2001; Liu et al., 2000). Due to the multifaceted nature of TCR engagement in live cells, the mechanism for the increase in functional avidity we report here is representative of *in vivo* antigen recognition and may be unrelated to alterations in the affinity between TCR and CD1b that could be measured using standard biophysical approaches (Cho et al., 2001).

Our data have important implications for antigen recognition *in vivo*. Previous studies have highlighted that subtle differences within the lipid tail can significantly influence TCR affinity for antigen, which translates into differences in function (Chancellor et al., 2017; de Jong et al., 2007; James et al., 2018; Van Rhijn et al., 2017). Our data suggest that co-receptor expression must also be factored into this calculation as our single cell analyses of SGL- and GMM-specific T cells demonstrate that glycolipid-specific T cells that express CD4 and CD8 co-receptors are functionally distinct. Indeed, our data suggest that these subsets may align closely with peptide-specific T cells, rather than iNKT or MAIT cells (James and Seshadri, 2020). Mycobacterial glycolipid-specific T cells that express the CD8 co-receptor are broadly cytotoxic T cells and express IFN-γ, TNF, and IL-2. This is consistent with the original description of an SGL-specific T cell clone, which produced IFN-g in the presence of M.tb infected cells and reduced the bacterial burden of M.tb in culture (Gilleron et al., 2004). Conversely, T cells that express the CD4 co-receptor are more diverse. They express some Th1 cytokines (TNF), but also can express features of Treg (FOXP3, CTLA4), Th2 (GATA3), and T_FH_ (BCL6) T cell subsets. At present, the only transcription factor that has been evaluated in CD1b-restricted T cells is PLZF, the transcription factor that regulates innate-like phenotypes in iNKT and MAIT cells (Constantinides et al., 2014; Martin et al., 2009; Savage et al., 2008). In a humanized transgenic mouse model, CD1b-restricted T cells did not express PLZF, which indicates that CD1b-restricted T cells may have a transcriptional landscape that is similar to MHC-restricted T cells (Zhao et al., 2015). Our findings that CD4 and CD8 CD1b-restricted T cells share major transcriptional pathways with MHC-restricted T cells is consistent with this model (James and Seshadri, 2020). However, we were unable to detect an enrichment of Runx3 in CD8 CD1b-restricted T cells, highlighting that there may also exist transcriptional pathways that are unique to CD1b-restricted T cells (Shan et al., 2017)

We detected a relationship between CD4 expression and Ki-67 expression, suggesting that the differences in functional avidity we describe here may be important for lipid antigen recognition during active TB in humans and the development of lipid-based vaccines. We noted a strong trend towards enrichment of activated T cells among CD1b-restricted T cells when compared to tetramer-negative T cells, highlighting that CD1b-restricted T cells appear to be actively proliferating in the context of active TB, likely as a result of antigen-specific activation. Consistent with this is the observation that only 1.8% of peripheral blood CD4 T cells express Ki67 during active TB (Adekambi et al., 2015). These findings suggest that CD4, and not CD8, T cells may be the major CD1b-restricted T cell subset activated during active TB, possibly as a result of enhanced functional avidity. Lipid-based vaccines have shown promise in pre-clinical animal models (Shang et al., 2018). Our data highlight the importance of considering whether CD4 or CD8 T cells would be targeted by the adjuvant or delivery platform when ‘tuning’ the chemical structure of the antigen to modify TCR avidity and immunogenicity of these novel vaccine strategies.

## MATERIALS AND METHODS

### Clinical Cohorts

For *ex vivo* analysis of SGL-CD1b tetramer positive cells, we studied two cohorts of healthy participants. First, U.S. healthy controls were recruited and enrolled at the Seattle HIV Vaccine Trials Unit as part of a cohort of healthy adults who to provided blood samples for developing and testing new assays. Peripheral blood mononuclear cells (PBMC) were collected by leukapheresis from five HIV-seronegative individuals with a known T cell response to CMV were used here. Second, we studied a subset of 6363 South African adolescents who were enrolled into a study that aimed to determine the incidence and prevalence of tuberculosis infection and disease in South Africa (Mahomed et al., 2011). Adolescents aged 12- to 18-years-old were enrolled at eleven high schools in the Worcester region of the Western Cape of South Africa. Participants were screened for the presence of latent tuberculosis infection (LTBI) by a tuberculin skin test and/or IFN-γ release assay (IGRA) QuantiFERON-TB GOLD In-Tube (QFTG) (Cellestis Inc.) at study entry. PBMC were isolated from freshly collected heparinized blood via density centrifugation and cryopreserved. For this work, a sample of five M.tb-infected (QFTG+) and five M.tb-uninfected (QFTG-) adolescents were selected based on availability of PBMC.

We also utilized cryopreserved PBMC samples isolated from adults with a new diagnosis of active tuberculosis also from the Worcester region of the Western Cape of South Africa (Day et al., 2011). Participants were over 18 years of age and HIV uninfected. All patients had positive sputum smear microscopy and/or were positive culture for M.tb. Blood was obtained and PBMC archived prior to or within 7 days of starting standard course anti-TB treatment, which was provided according to guidelines of the South African National Tuberculosis Programme.

### Ethics Statement

The study protocols were approved by the IRBs of the University of Washington, the Fred Hutchinson Cancer Research Center, and the University of Cape Town. Written informed consent was obtained from all adult participants as well as from the parents and/or legal guardians of the adolescents who participated. In addition, written informed assent was obtained from the adolescents.

### Culture Media

Media (R10) for washing PBMC consisted of RPMI 1640 (Gibco, Waltham, MA) supplemented with 10% fetal calf serum (Hyclone, Logan, UT). Our base T cell media (TCM) consisted of sterile-filtered RPMI 1640 supplemented with 10% fetal calf serum, 100 U/ml Penicillin, 100 mg/ml Streptomycin, 55 mM 2-mercaptoethanol, 0.3X Essential Amino Acids, 60 mM Non-essential Amino Acids, 11 mM HEPES, and 800 mM L-Glutamine (Gibco, Waltham, MA). Our TCM containing human serum (TCM/HS) consisted of sterile-filtered RPMI 1640 supplemented with 10% human serum (derived from healthy donors), 100 U/ml Penicillin, 100 mg/ml Streptomycin, and 400 mM L-Glutamine (Gibco, Waltham, MA). For culture of Jurkat cells, enhanced RPMI was used, which consisted of RPMI 1640 (Gibco, Waltham, MA) supplemented with 10% fetal calf serum (Hyclone, Logan, UT), 100 U/ml Penicillin, 100 mg/ml Streptomycin, and 800 mM L-Glutamine (Gibco, Waltham, MA).

### Generation of SGL- and GMM-loaded Tetramers

Soluble biotinylated CD1b monomers were provided by the National Institutes of Health Tetramer Core Facility (Emory University, Atlanta, GA). The loading protocol for CD1b monomers was based on previously published loading protocols (James et al., 2018; Kasmar et al., 2011). For SGL-loaded CD1b tetramers, a biologically validated synthetic analog of SGL, referred to as AM Ac2SGL in James et al., 2018, was used for all experiments except those involving single-cell profiling for which we used SGL purified from cell wall extracts from *Mycobacterium tuberculosis* (Gilleron et al., 2004; James et al., 2018). SGL was dried down in a glass tube in a stream of nitrogen and sonicated into a 50 mM sodium citrate buffer at pH 4, containing 0.25% 3-[(3-cholamidopropyl)dimethylammonio]-1-propanesulfonate (CHAPS) (Sigma, St. Louis, MO) for two minutes at 37°C. For GMM-loaded CD1b tetramers, C32 GMM derived from *Rhodococcus equi* was dried down and sonicated into 50 mM sodium citrate buffer at pH 4, containing 0.25% CHAPS. The sonicate was transferred to a microfuge tube, and 20 μl of CD1b monomer was added and incubated in a 37°C water bath for 2 hours with vortexing every 30 minutes. At the end of the incubation, the solution was neutralized to pH 7.4 with 6 μl of 1M Tris pH 9. After addition of CD1b, the mixture was incubated in a 37°C water bath for 2 hours. Finally, 10 μl of Streptavidin conjugated to ECD or APC (Life Technologies, Carlsbad, CA) was added in ten aliquots of 1 μl every 10 minutes. The final product was filtered through a SpinX column (Sigma, St. Louis, MO) to remove aggregates and stored at 4°C until use.

### Tetramer Staining and Sorting

#### Ex vivo analysis

PBMC were thawed in warm thaw media (R10 with 2 μl/ml Benzonase (Millipore, Billerica, MA) sterile-filtered) and centrifuged at 1500 rpm for 5 minutes. The supernatant was decanted, and the cells were resuspended in R10 and counted by Trypan Blue (Millipore, Billerica, MA) exclusion. The cells were centrifuged at 1500 rpm for 5 minutes and plated at a density of 1 million cells per well in a 96-well U-bottom plate. A portion of the PBMC were resuspended in R10 at a density of 2 million cells per ml in 50 ml conicals with the caps lightly in place and rested overnight at 37°C in humidified incubators supplemented with 5% CO2. The PBMC in the 96-well plate were washed with FACS buffer (1x phosphate-buffered saline (PBS) (Gibco, Waltham, MA) supplemented with 0.2% bovine serum albumin (BSA) (Sigma, St. Louis, MO) and centrifuged at 1800 rpm for 3 minutes. Next, the cells were washed twice with PBS and stained with Aqua Live/Dead stain (Life Technologies, Carlsbad, CA) according to the manufacturer’s instructions. Following a 15 minute incubation at room temperature, the cells were washed twice in PBS. They were then blocked with human serum (Valley Biomedical, Winchester, VA) and FACS buffer mixed 1:1 for 10 minutes at 4°C. The wells were washed twice with FACS buffer and then resuspended in 50 μl FACS buffer with 1 μl of unloaded CD1b tetramer and 1 μl of each SGL-loaded CD1b tetramer labeled with APC or ECD and incubated at room temperature for 40 minutes in the dark. After this incubation period, the cells were washed twice with FACS buffer and then labelled with anti-CD3 ECD (Beckman Coulter, Brea, CA), anti-CD4 APC Ax750 (Beckman Coulter, Brea, CA), and anti-CD8ß BB700 (BD Biosciences, San Jose, CA) antibodies for 30 minutes at 4°C. After two final washes in FACS buffer, the cells were fixed in 1% paraformaldehyde (PFA) (Electron Microscopy Sciences, Hatfield, PA) and acquired on a BD LSRFortessa (BD Biosciences, San Jose, CA) equipped with blue (488 nm), green (532 nm), red (628 nm), violet (405 nm), and ultraviolet (355 nm) lasers.

#### In vitro-derived T cell lines and transduced T cells

T cell lines or transductants were plated at one million cells per well in a 96-well U-bottom plate. Cells were washed twice with PBS and resuspended with Live/Dead Fixable Aqua or with Live/Dead Fixable Green Dead Cell Stain Kit (Life Technologies, Carlsbad, CA) per the manufacturer’s instructions. For this step and all subsequent steps, the cells were kept in the dark. Following a 15 minute incubation, cells were washed twice with PBS and blocked with human serum (Valley Biomedical, Winchester, VA) prepared in FACS buffer (1x PBS (Gibco, Waltham, MA) supplemented with 0.2% BSA (Sigma, St. Louis, MO)) mixed 1:1 for 10 minutes at 4°C. Cells were then resuspended in 50 μl of FACS buffer containing 1 μl of SGL-loaded CD1b and 1 μl of mock-loaded control CD1b tetramers, then incubated at room temperature for 60 minutes. The tetramer titers were determined prior to use in the present study (data not shown). Following a 15 minute incubation at room temperature, the cells were washed twice in PBS and then stained with anti-CD3 ECD (Beckman Coulter, Brea, CA), CD4 APC Ax750 (Beckman Coulter, Brea, CA) and anti-CD8α PerCP Cy5.5 (BD Biosciences, San Jose, CA). When T cells transduced with exogenous TCR were used, anti-mouse TCR β chain APC (BD Biosciences, San Jose, CA) was also included in the antibody cocktail. The optimal titers of all antibodies were determined prior to use (data not shown). After two final washes in FACS buffer, the cells were fixed in 1% paraformaldehyde (PFA) (Electron Microscopy Sciences, Hatfield, PA) and acquired on a BD LSRFortessa as above.

#### Sorting

The transduced T cells were sorted to purity using a modified version of the tetramer staining method described above. However, after the antibody stain, the transduced T cells were resuspended in 200 μl of FACS buffer and tetramer-positive T cells were sorted at the UW Department of Immunology Flow Cytometry Core using a FACS Aria II (BD Biosciences, San Jose, CA) cell sorter equipped with blue (488 nm), red (641 nm), and violet (407 nm), lasers. Cells were sorted into 3 ml of TCM in 4 ml FACS tubes.

### T cell Receptor Cloning and Transduction

#### Template-switched PCR

The sequences of the A01 and A05 TCR were determined by a previously described 5’-rapid amplification of cDNA ends (5’RACE) based cloning strategy based on the manufacturer’s instructions (Yu et al., 2018; Jing et al., 2017) (Takara Bio, Japan). Briefly, RNA was extracted from T cells and cDNA was synthesized using a dT oligo and template-switch adaptor. Then, customized oligos targeting the TCRa and TCRß constant region were used to amplify TCR sequence when combined with a universal primer. The oligo sequences are as follows:

TCRα chain primer: 5’-GATTACGCCAAGCTTGTTGCTCCAGGCCACAGCACTGTTGCTC-3’
TCRβ chain primer: 5’-GATTACGCCAAGCTTCCCATTCACCCACCAGCTCAGCTCCACG.3’ The products were cloned into sequencing vector and sequenced using Sanger sequencing. The variable genes, joining genes, and CDR3 used by the sequences were analyzed using the IMGT Database (Montpellier, France).

#### T cell Receptor Cassette Construction

Codon-optimized A01 and A05 T cell TCR sequences were assembled into a TCR cassette (Linnemann et al. 2013). In this construct, the TCR α and TCR β chain constant regions are replaced with modified murine TCR α chain and TCR β chain constant regions (mTCRBC) to facilitate measurement of TCR expression by flow cytometry. It also contains an extra cysteine residue to encourage pairing between exogenous TCR chains. The TCR cassettes were synthesized through Thermo Fisher GeneArt Synthesis service and were then cloned into pRRL.PPT.MP.GFPpre (Jing et al., 2016; Zhou et al., 2012) using BamHI and SalI restriction enzymes (New England BioLabs, Ipswich, MA). All plasmids were purified using Maxi Prep kits (Qiagen, Hilden, Germany). GenBank accession number for the A01 and A05 TCR constructs is pending.

#### Generation of Lentivirus

Lenti-X HEK293T cells (Clontech, Mountain View, CA) were seeded at 2 million cells per 100 mm tissue culture dish and incubated for 48 hours at 37°C/5% CO2 in DMEM (Gibco, Waltham, MA) or until cells reached 75% confluency. The medium was replaced 4 hours before transfection. Cells were transfected with 10 μg pRRL-TCR plasmid, 5 μg pCI-VSVG envelope plasmid, and 5 μg of a psPAX2 packaging vector (gifted from Dr. Stanley Riddell at Fred Hutchinson Cancer Research Center). Plasmids were mixed with Fugene 6 transfection reagent (Promega, Madison, WI) at a dilution of 1:12 in a total volume of 600 μl. Transfection mixture was added dropwise into the cell culture and incubated overnight in the conditions described above. The medium was then replaced and incubated for an additional 48 hours. After this time, 20 μl of supernatant was titered using Lenti-X GoStix (Clontech, Mountain View, CA) per the manufacturer. Supernatant was then harvested every 12 hours for a total of three collections. At each collection, cell debris was removed by centrifugation at 1500 rpm for 5 minutes, and cleaned supernatant was reserved in a 50-ml conical and kept at 4°C until three collections had been acquired. Supernatant was then incubated overnight with Lenti-X concentrator (Clontech, Mountain View, CA) at a ratio of 1:3. The following day, the supernatant was centrifuged at 1500g for 45 minutes at 4°C. Supernatant was then discarded, and the pelleted virions were resuspended in 300 μl R10 media and stored at −80°C until further use. Viral preparations were titred for optimal transduction efficiency in Jurkat cells.

#### Transduction of Jurkat Cells

CD8-expressing and DN Jurkat cells were seeded at 1 million cells per well in a 48-well plate in enhanced RPMI. Jurkat cells were generously provided by Dr. Thomas Blankenstein. The same day, the Jurkat cells were transduced with TCR lentivirus at an estimated multiplicity of infection (MOI) of 5 with 1 μl of polybrene at a final concentration of 4 μg/ml (Sigma, St. Louis, MO). Cells and virus were incubated for 4 hours at 37°C/5% CO2 and washed with PBS (Gibco, Waltham, MA) to remove excess virus. Jurkat cells were maintained in culture for 1 week using enhanced RPMI and screened for TCR expression (Jing et al., 2016) using an anti-mouse TCR β chain (mTCRBC) APC antibody (BD Biosciences, San Jose, CA) and tetramer staining as described above.

#### Transduction of Primary T cells

Cryopreserved PBMC from an anonymous blood bank donor (Bloodworks Inc, Seattle, WA) was thawed as described above and enumerated using trypan blue exclusion. T cells were isolated from PBMC using magnet activated cell sorting (MACS) using the Pan T cell Isolation Kit (negative selection) according to the manufacturer’s instructions (Miltenyi Biotec, Germany). Following separation, T cells are enumerated using Trypan Blue exclusion and co-incubated with CD3/CD28 Human T cell Activating Dynabeads (ThermoFisher) at a 3:1 bead:T cell ratio in a total volume of 1 ml TCM supplemented with 50 units/ml recombinant IL-2 in a 48-well plate. On day 2, lentiviral stock and 4 μg/ml polybrene was added to culture. Viral titer is determined prior to use (data not shown). Following viral addition to the well, the 48-well plate is then centrifuged at 679 x g for 90 minutes at 32°C. The cells are then incubated at 37°C/5% CO2 for 48 hours. On day 4, the cells are harvested and transferred to a 4 ml FACS tube and rinsed with 3 ml of TCM. The cells are then centrifuged at 1,300 rpm for 5 minutes to remove the polybrene from the cells. Following centrifugation, the cells are resuspended in TCM supplemented with rIL-2 as described above and returned to the original well. The cells are then incubated for 72 hours at 37°C/5% CO2. On day 7, the cells are harvested and moved to a 4ml FACS tube and incubated on a magnetic stand for 10 minutes at room temperature to remove the Dynabeads. The media, which contains the activated T cells, was then removed and added to a 15 ml conical. Fresh media is then added to the 4 ml FACS tube to rinse the beads, and the 4 ml FACS tube is then returned to the magnetic stand and incubated for 5 minutes at room temperature. The media is then removed from the 4 ml FACS tube and added to the 15 ml conical. Cells are then centrifuged at 1,300 rpm for 5 minutes, and resuspended in fresh TCM supplemented with rIL-2 and returned to the 48-well plate. Cells are then incubated for 4 days at 37°C/5% CO2 and maintained as necessary. On day 12, the cells are then assayed to quantify T cell transduction by tetramer staining as described above.

### T cell Culture and Analysis

#### Rapid Expansion Method

Further expansion of the T cells transduced with exogenous TCR was performed using a modified version of a previously established rapid expansion protocol (Riddell et al., 1992). Briefly, 100,000 T cells were mixed with 5 million irradiated EBV-transformed B cells and 25 million irradiated PBMC as feeder cells in R10 in T25 tissue culture flasks (Costar, St. Louis, MO) with 25 ml TCM. Anti-CD3 (clone OKT3) was added at a final concentration of 30 ng/ml, and the mixture was incubated overnight at 37°C/5% CO2. The following day, recombinant IL-2 (rIL-2) (Prometheus Pharmaceuticals through UWMC Clinical Pharmacy) was added at a final concentration of 50 U/ml. On day 4, the cells were washed twice in TCM to remove the anti-CD3 antibody and resuspended in fresh media supplemented with rIL-2 at 50 U/ml. Half the media was replaced every three days or split into new T25 tissue culture flasks as determined by cell confluency. After 13 days in culture, the transduced T cells were screened by tetramer staining and then frozen on day 14.

#### Limiting Dilution Cloning

Sorted T cells were washed and resuspended in TCM/HS and plated at 1 or 2 cells per well in each well of a 96-well plate to create single cell clones. Irradiated PBMC (150,000 cells per well) were added as feeder cells along with PHA (Remel, San Diego, CA) at a final concentration of 1.6 μg/ml. After two days in culture at 37°C/5% CO2, 10 μl natural IL-2 (Hemagen, Columbia, MD) was added to each well. Half the media was replaced every two days with TCM/HS and natural IL-2. Cultures were maintained for 14 days and screened for antigen specificity by tetramer staining and IFN-γ ELISPOT.

#### Intracellular Cytokine Staining

On day 0, cryopreserved T cell lines were thawed, counted, and enumerated using Trypan blue exclusion T cell lines were rested in TCM overnight as described above and enumerated again on day 1. On day 0, SGL was evaporated to dryness from chloroform-based solvents under a sterile nitrogen stream and then sonicated into media. This lipid suspension was added to 50,000 K562-EV or K562-CD1b cells at 5 mg/ml final concentrations. K562 cells were incubated for 18 hours at 37°C, 5% CO2 to facilitate lipid loading. On day 1, rested T cell lines were split and added to the K562 cells without washing at a final density of 1 million/well. Thus, each T cell line was co-incubated with loaded K562-CD1b and K562-EV antigen presenting cells. The cell mixture was allowed to incubate for 6 hours in the presence of anti-CD28/Ly29d Abs (BD Biosciences, San Jose, CA), brefeldin A at a final concentration of 10 mg/ml (Sigma-Aldrich), and GolgiStop containing Monensin (BD Biosciences, San Jose, CA), after which EDTA, at a final concentration of 2 mmol, was added to disaggregate cells. On day 2, the samples were washed twice in PBS and then stained with Aqua Live/Dead (Life Technologies, Carlsbad, CA) prepared according to manufacturer’s instructions and incubated for 20 minutes at room temperature. Live/Dead staining and all steps following were performed in the dark. The cells were washed twice in PBS and then incubated at room temperature for 10 minutes in 1x FACS Lyse (BD Biosciences, San Jose, CA). Following one wash with FACS buffer, the cells were incubated an additional 10 minutes in 1x FACS Perm II (BD Biosciences, San Jose, CA) at room temperature. The cells were washed twice in FACS buffer and labeled with antibodies for CD3, CD4, CD8, IFN-γ, IL-2, IL-4, CD154 (BD Biosciences, San Jose, CA), and TNF (eBioSciences, Waltham, MA) for 30 minutes at 4°C. Following two final washes in FACS buffer, the cells were fixed in 1% PFA and acquired on a BD LSRFortessa as above (BD Biosciences, San Jose, CA).

#### IFN-γ and Granzyme B ELISPOT

EMD Multiscreen-IP filter plates (Millipore, Billerica, MA) were coated with 1D1K antibody (Mabtech, Sweden) diluted 1:400 in PBS and incubated overnight at 4°C. The following day, one thousand T cells were plated 1:50 with K562-CD1b and K562-EV cells. Lipids antigens were stored in chloroform:methanol (2:1, v:v) at a stock concentration of 1 mg/ml and a temperature of - 20°C. For these experiments, approximately 4 μl were dried in glass centrifuge tubes under a sterile nitrogen stream. The lipids were sonicated into TCM to obtain a 4 μg/ml suspension and plated with the cells at a final concentration of 1 μg/ml. These cultures were incubated at 37°C for approximately 16 hours. The following day, the cells were washed twice with sterile water to lyse the cells, and the plates were incubated with the detection antibody 7-B6-1-biotin (Mabtech, Sweden) diluted 1:1000 in PBS + 0.5% FCS and incubated for two hours at room temperature. The cells were then washed five times with PBS and incubated in ExtrAvidin-Alkaline Phosphatase (Sigma, St. Louis, MO) diluted 1:1000 in PBS and incubated for one hour at room temperature. The wells were then washed five times and incubated with BCIP/NBT substrate (Sigma, St. Louis, MO) for five minutes to develop the membrane. The wells and IFN-γ spots were counted using an ImmunoSpot S6 Core Analyzer (Cellular Technology Limited, Cleveland, OH). The granzyme B ELISPOT followed identical methods with the following exceptions: EMD filter plates were coated with GB10 antibody diluted to 15 μg/ml, GB11-biotin diluted to 1 μg/ml was used as the detection antibody, and the culture incubation was extended to 48 hours (Mabtech, Sweden).

#### Cytotoxicity Assay

To detect cytotoxic activity, the T cell lines were thawed and rested overnight in TCM as described above. The following day, K562-CD1b and K562-EV were stained with Cell Trace Violet (Invitrogen, Carlsbad, CA) at 10 μM and 0.2 μM, respectively, for 20 minutes at 37°C. Staining was quenched by adding 10 ml cold PBS to the staining solution. Stained K562-CD1b and K562-EV cells were then mixed at a 1:1 ratio. This cell mixture was then co-incubated with the T cell lines at an effector to target ratio of 5:1 in a 96-well plate in a total volume of 200 μl of TCM. SGL was added to the culture media at 1 μg/ml, and the cell suspension was centrifuged briefly to pellet the cells. Cells were incubated for 24 hours at 37°C/5% CO2. The following day, the cells were washed twice with PBS, and stained with Aqua Live/Dead (Life Technologies, Carlsbad, CA) prepared according to manufacturer’s instructions and incubated for 20 minutes at room temperature. The cells were then washed twice with PBS and then stained with anti-CD3 antibody (BD Biosciences, San Jose, CA) for 30 minutes at 4°C in FACS buffer. The cells were then washed twice with FACS buffer and fixed in 1% PFA. Samples were acquired on an LSR Fortessa as above (BD Biosciences, San Jose, CA). We defined “% Cell Death” as the percent change of the proportion of K562-CD1b cells that remain after co-culture compared to the proportion of K562-CD1b cells at baseline. The “baseline” condition is defined by the mixture of K562-CD1b and K562-EV cells where no T cells or lipid antigen was present in the culture (Supplemental Figure 8).

### Single-cell Transcriptional Profiling

Transcriptional profiles and T cell receptor sequences were determined from sorted cells using an established protocol (Han et al., 2014). Briefly, cells were sorted from cryopreserved PBMC and T cell lines as positive controls using CD1b tetramers loaded with GMM or purified SGL into single wells of a 96-well PCR plate (Eppendorf, Hamburg, Germany), which contained 5 μl of sort buffer consisting of (1X Qiagen One-step RT PCR Buffer (Qiagen, Hilden, Germany), 0.1 mM dithiothreitol (Invitrogen, Carlsbad, CA), RNaseOUT (Invitrogen, Carlsbad, CA), and molecular-grade nuclease-free water (Fisher, Hampton, NH)). Cells were centrifuged briefly at 1800 rpm for 1 minute, and frozen at −80°C overnight. The following day, the reverse transcription reaction was performed using primer pools containing TCRAV and TCRBV primers at 0.06 μM final concentration, TCRA and TCRB constant region primers at 0.3 μM final, and phenotype gene primers at 0.1 μM. Primer sequences are as described in Han et al., 2014, with exceptions as noted in Supplemental Tables 5 and 6. Samples then undergo nested amplification in a Mastercycler Nexus Gradient Thermal Cycler (Eppendorf, Hamburg, Germany). The second phase of amplification uses 0.4 units of HotStarTaq DNA polymerase (Qiagen, Hilden, Germany). Lastly, the amplicons were barcoded according to their plate, row, and column location using the barcodes specified in the published protocol (Han et al., 2014). The TCR α chain, TCR β chain, and phenotyping genes were amplified separately at this step. The row and column barcodes were each added at 0.05 μM, and Illumina MiSeq paired end adapters were used at 0.5 μM. After the amplification steps, all wells are pooled from each plated and cleaned using Ampure XP PCR Cleanup Beads (Beckman Coulter, Brea, CA) per the manufacturer’s instructions. Each plate was pooled and the plate libraries were quantified on a Qubit Fluorometer 3.0 using a dsDNA High-sensitivity Kit (Life Technologies, Carlsbad, CA). The plates were then pooled in equal volumes and the pooled library was then diluted to 6 μM and sequenced using a 500-cycle V3 reagent on an Illumina MiSeq (Illumina, San Diego, CA), which yields 25 million paired reads of 250 base pairs, at the Fred Hutchinson Cancer Research Center Genomics Core.

### Single-cell Computational Methods

Raw reads were trimmed and demultiplexed using VDJFasta software (Glanville et al., 2009). Paired end reads were assembled by finding a consensus of at least 100 bases. Amplicons smaller than 100 bases were removed. The resulting paired-end reads were then assigned to wells according to barcodes that designate the plate, row, and column. The average read count for this study was approximately 47,000 reads per well.

To map the TCR reads, a cutoff of >95% sequence identity was established to collapse reads and establish a consensus sequence within each well. All sequences exceeding 95% sequence identity are assumed to derive from the same TCR sequence. The 95% cutoff conservatively ensures all sequences derived from the same transcript would be properly assigned. In addition, TCR assignments that fail to pass a minimum frequency of 0.1 for each well are removed from the dataset as they might be derived from sorted doublets (Han et al., 2014). For this dataset, the median frequency of the dominant TCR frequency in our dataset is 0.79 with a range of 0.313-0.999.

For phenotyping transcripts, the number of reads containing a 95% match to the customized database of transcription factor and cytokine genes are scored. Genes profiled in this study are summarized in Supplemental Tables 5 and 6. The resulting data is then compiled into a .csv file containing the TCR assignment and number of reads for each phenotype gene for each well. A minimum read count of five reads per gene was used to define expression of each gene as established in Han et al., 2014.

Data were then input into the R programming environment and further processed and formatted. The resulting matrix was then analyzed using Seurat to cluster the data and perform feature selection for two analyses (Satija et al., 2015). The first was unsupervised and included all cells that passed our QC criteria (n = 272). The second was a supervised analysis that only included cells where either CD4 or CD8 was detected (n = 105). The proportion of cells from the tetramer-positive and tetramer-negative groups that were included in the final analysis were similar (39% and 37%, respectively). Annotated data, as well as the code used to generate these data and figures are available at: https://github.com/chetanseshadri. Other data visualization and statistical tests were performed in Graph Pad Prism (v6) (San Diego, CA).

### Computational and Statistical Methods

Initial compensation, gating, and quality assessment of flow cytometry data was performed using FlowJo version 9 (FlowJo, TreeStar Inc, Ashland OR). Flow cytometry data processing and MFI extraction was performed using the OpenCyto framework in the R programming environment (Finak et al., 2014). Statistical tests are described in the Figure and Table legends. Categorical variables were analyzed using a Fisher’s exact test. When two continuous variables were analyzed, a Student’s t-test was used when data were normally distributed, and a Mann-Whitney test was used when normality assumption could not be met. When more than two continuous variables were analyzed, a two-way analysis of variance (ANOVA) was used when data were normally distributed, and a Kruskal-Wallis test was performed when a normal distribution could not be assumed. Post-hoc Dunn tests were performed after ANOVA or Kruskal-Wallis tests to determine which group(s) were different. A Wilcoxon Signed Rank test was used when the continuous variables were not independent. When multiple hypotheses were tested, p-values were adjusted using the Bonferroni method. These tests were conducted in R (v3.8.5) or Graph Pad Prism version 6 (GraphPad, GSL Biotech, San Diego, CA).

## ACKNOWLEDGEMENTS

The authors thank Krystle Yu, Paula Marsland, Natalie Erjavec, Sarah Roepke, and Martin Prlic for their technical assistance, D. Branch Moody for supplying purified GMM for these studies, and Stephen De Rosa for supplying the Seattle Assay Control samples. This work was supported by the U.S. National Institutes of Health (R01AI12518904 to CS) and the Doris Duke Charitable Foundation (Grant No. 2016103 to CS). CAJ was supported by the Institute of Translational Health Sciences (5TL1TR00231803) and the Molecular Medicine Training Program (T32GM095421).

## SUPPLEMENTARY MATERIAL

**Supplemental Table 1.**
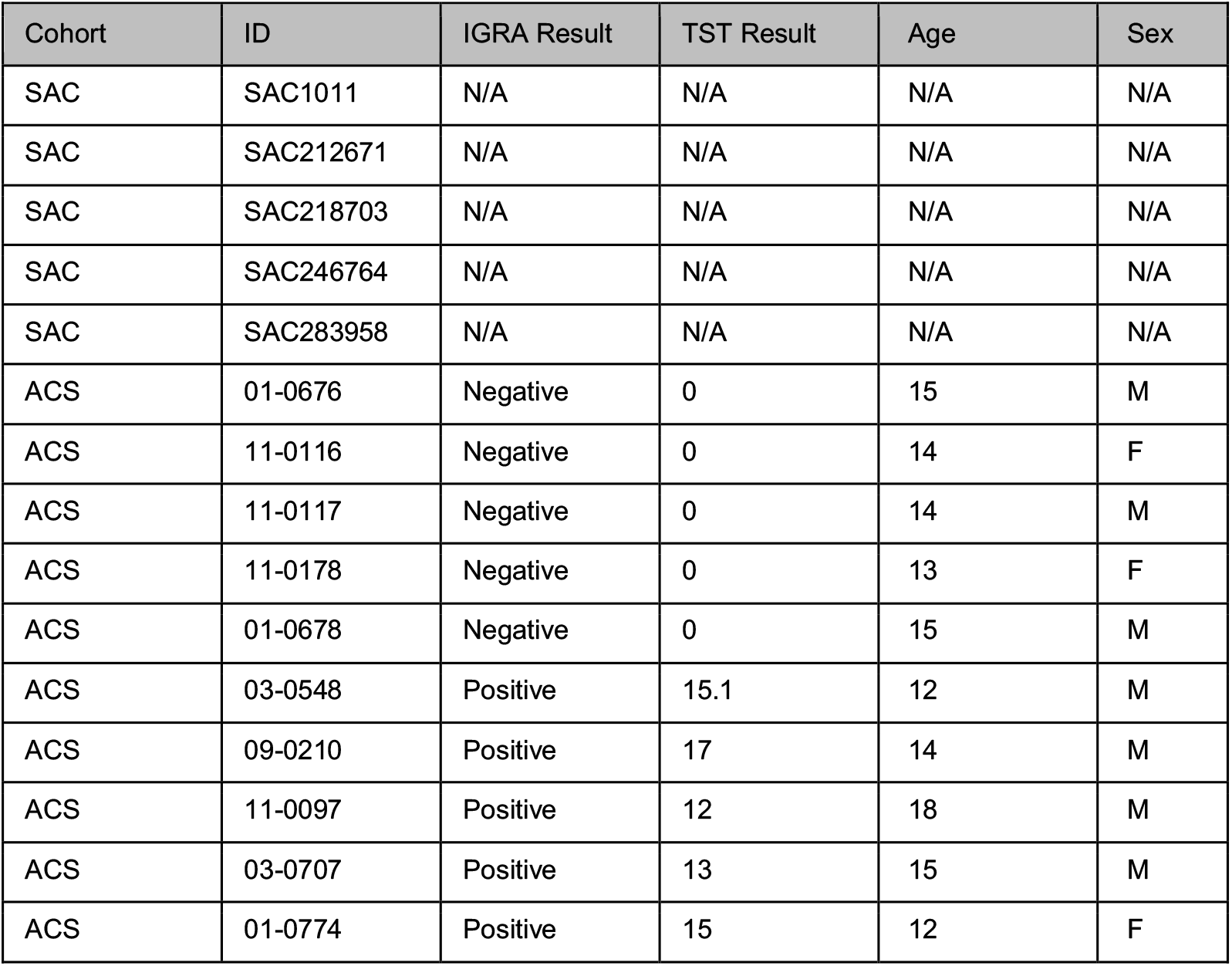
Human samples used for *ex vivo* analysis of SGL-specific T cells. Human samples used in this study are listed by cohort of origin and participant identifier (ID). Seattle Assay Controls (SAC) are healthy donors from Seattle, WA, that were enrolled through the HIV Vaccine Trials Network (HVTN). The Adolescent Cohort Study (ACS) is composed of M.tb-infected or M.tb-uninfected adolescents from South Africa. Interferon-γ release assay (IGRA) and tuberculin skin test (TST) results are listed and concordantly positive or negative (Mahomed et al., 2011). TST induration is reported here in mm. Age reflects the age at last birthday at the time the sample was collected. Sex is self-reported.

**Supplemental Table 2.** Flow cytometry panel used for *ex vivo* identification of SGL-specific T cells.

| Specificity | Purpose | Fluorochrome | Clone | Supplier |
| --- | --- | --- | --- | --- |
| CD3 | Lineage | BUV395 | UCHT1 | BD Biosciences |
| CD4 | Lineage | APC H7 | 13B8.2 | BD Biosciences |
| CD8β | Lineage | BB700 | 2ST8.5H7 | BD Biosciences |
| CD45RA | Memory | BUV737 | HI100 | BD Biosciences |
| CCR7 | Memory | BV711 | 150503 | BD Biosciences |
| Pan-$\gamma\delta$ | $\gamma\delta$ T cells | PE-Vio770 | 11F2 | Miltenyi Biotec |
| Vδ2 | Vγ9δ2 T cells | AF700 | B6 | BioLegend |
| TRAV1-2 | TCR Identification | BV605 | 3C10 | BioLegend |
| CD14 | Exclusion | V785 | M5E2 | BioLegend |
| CD19 | Exclusion | V785 | SJ25C1 | BioLegend |
| Fixable Green<br>Dead Cell Stain | Viability | FITC | N/A | Life Technologies |
| CD1b-SGL | M.tb lipid antigen-specific T cells | ECD | N/A | Custom |
| CD1b-SGL | M.tb lipid antigen-specific T cells | PE | N/A | Custom |
| CD1b-Mock | Exclusion | V510 | N/A | Custom |

**Supplemental Table 3.** Enrichment of V Genes Among SGL-CD1b and GMM-CD1b tetramer-sorted cells. The count of recovered TCRs using the specified V gene is reported from both patients studied, from GMM-CD1b or SGL-CD1b tetramer-sorted cells (TCRs from GMM-CD1b sorted cells = 96, TCRs from SGL-CD1b sorted cells = 127). IMGT nomenclature is used for variable (V) gene identification. Bulk T cell counts are reported as the sum of the templates from both patients that use the specified V gene using previously published ImmunoSEQ data from the same patients studied here (DeWitt et al., 2018). Template counts were used to populate a 2×2 contingency table and unadjusted p-values resulting from a Fisher’s exact test are shown.

| V Gene | Antigen | Tetramer-Positive |  | Bulk T cells |  | P-value |
| --- | --- | --- | --- | --- | --- | --- |
|  |  | Positive | Negative | Positive | Negative |  |
| TRAV1-2 | GMM | 10 | 86 | 7438 | 249848 | 0.0005 |
| TRAV8-6 | SGL | 6 | 121 | 5428 | 251858 | 0.053 |
| TRAV13-2 | SGL | 2 | 125 | 2514 | 254772 | 0.35 |
| TRAV21 | SGL | 8 | 119 | 7394 | 249892 | 0.031 |
| TRAV19 | SGL | 8 | 119 | 10745 | 246541 | 0.26 |

**Supplemental Table 4.** Differentially expressed genes among clusters visualized by UMAP. Clusters were defined and visualized as described in Figure 5 of the main text. The average log-fold-change indicates the difference in average expression between the cells within a cluster and outside the cluster. Positive values indicate that the gene is more highly expressed within the cluster, and negative values indicate that the gene is more highly expressed outside the cluster. P-values were calculated between the two groups of cells using the Wilcoxon Rank Sum Test and adjusted using Bonferroni correction using all the genes in the data set (n=23).

| Feature | Cluster | Average log-Fold Change | Adjusted p-value |
| --- | --- | --- | --- |
| PRF1 | 0 | -0.79 | 3.77e-11 |
| TBET | 0 | -0.64 | 2.40e-10 |
| EOMES | 0 | -0.70 | 9.58e-09 |
| CD8 | 0 | -0.33 | 2.33e-03 |
| GATA3 | 0 | -0.30 | 3.75e-03 |
| IFNG | 0 | -0.38 | 6.18e-03 |
| BCL6 | 0 | -0.30 | 9.58e-02 |
| CD4 | 1 | -0.69 | 3.73e-10 |
| TBET | 1 | 0.45 | 1.41e-06 |
| CD8 | 1 | 0.43 | 2.56e-06 |
| IFNG | 1 | 0.47 | 2.98e-05 |
| EOMES | 1 | 0.45 | 9.20e-05 |
| PRF1 | 1 | 0.44 | 1.39e-04 |
| BCL6 | 1 | -0.34 | 4.97e-02 |
| BCL6 | 2 | 0.70 | 2.11e-10 |
| MKI67 | 2 | 0.59 | 1.12e-06 |
| TNF | 2 | 0.48 | 7.22e-05 |
| CTLA4 | 2 | 0.49 | 2.11e-04 |
| FOXP3 | 2 | 0.37 | 2.77e-03 |
| CD4 | 2 | 0.31 | 5.02e-03 |
| IL12A | 2 | 0.25 | 2.34e-02 |
| GATA3 | 2 | 0.29 | 4.71e-02 |

**Supplemental Table 5.**
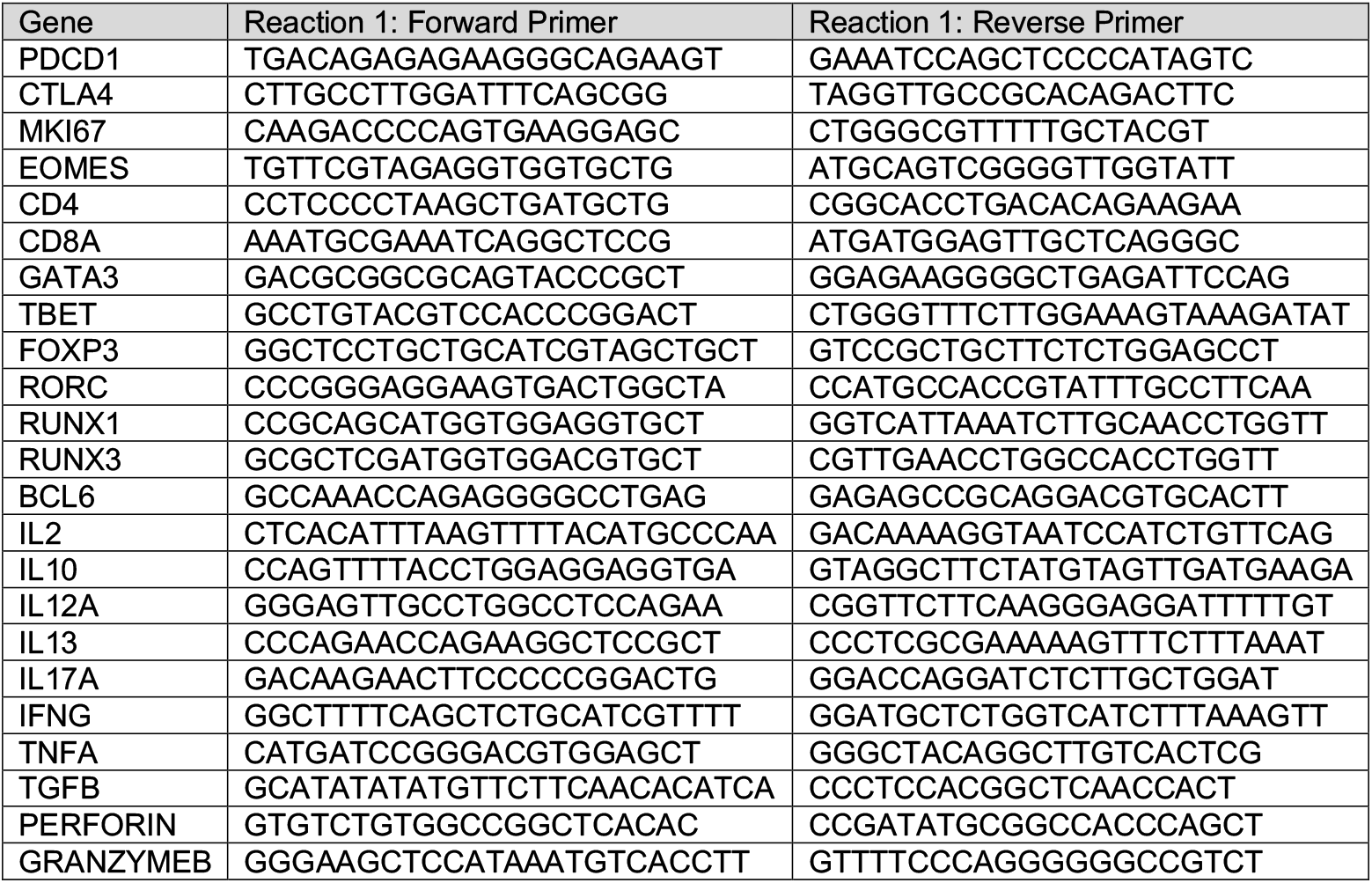
Gene-specific primers used for transcriptional profiling reaction 1.

**Supplemental Table 6.**
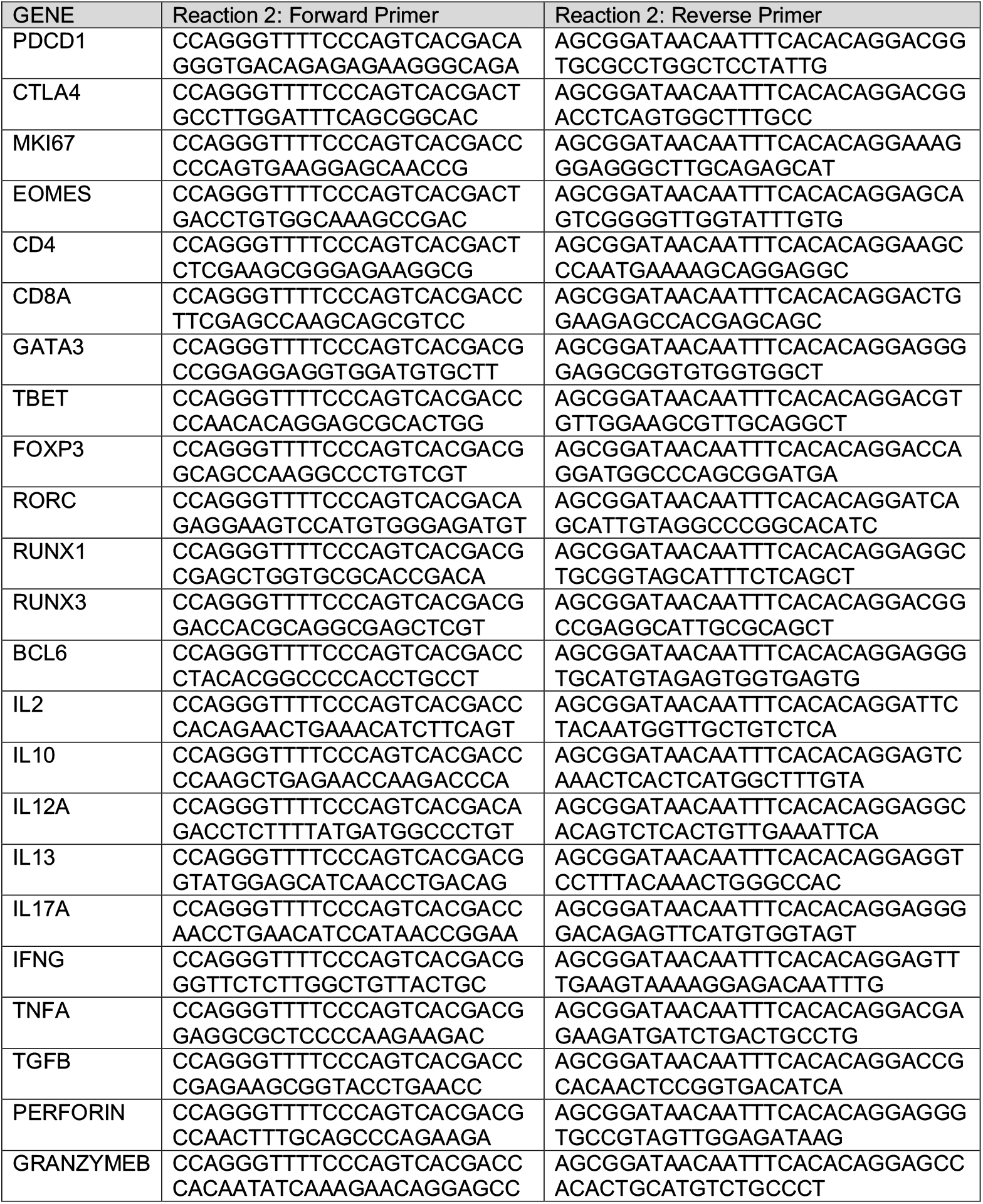
Gene-specific primers used for transcriptional profiling reaction 2.

## SUPPLEMENTAL FIGURE LEGENDS

**Supplemental Figure 1. Gating Strategy for Flow Cytometry and Cell Sorting. (A)** Representative gating strategy for identification of SGL-specific T cells. The gates proceeded from Live and CD3^+^ cells to CD14^-^ and CD19^-^ cells to single cells to a CCR7 “keeper gate” to lymphocytes by size gating. The CCR7 Keeper Gate is applied to eliminate events stained with dye aggregates, as these can interfere with downstream analysis and gating of rare cell populations (Layton et al., 2018). After this point, gates were drawn for γδ T cells, TRAV1-2, CD4 and CD8, and CD45RA and CCR7 independently for CD4 and CD8 T cells. (B) Gates were drawn for SGL-CD1b tetramer-positive cells as defined by dual staining with two SGL-CD1b tetramers, and negative for Mock-loaded CD1b tetramer. (C) Boxplots depict the median and interquartile range of SGL-CD1b frequency, expressed as a percentage of CD3^+^ T cells. Each dot represents the percentage of SGL-CD1b-specific T cells from one donor. (Kruskal Wallis with Dunn posttest, p = 0.4, n = 15). (D) Boxplots depict the median and interquartile range of CD3 MFI of all T cells within each co-receptor group (double positive (DP), CD4, CD8, and double negative (DN)). Each dot represents the percent of one sample. The percent of cells in each group was compared to that present in total CD3+ T cells (grey). (Kruskal Wallis with Dunn post-test, *** = p < 0.0001, * = 0.001 < p < 0.01, n = 15). (E) Natural SGL-CD1b and GMM-CD1b tetramers were incorporated into a multi-parameter flow cytometry assay to isolate SGL-specific and GMM-specific T cells using fluorescence activated cell sorting (FACS). The tetramer positive gate was defined by a Mock-loaded CD1b negative control tetramer (left) and a positive control using SGL- and GMM-specific T cell lines diluted in donor PBMC (middle, right). (F) Natural SGL-CD1b and GMM-CD1b tetramer positive T cells in the blood were sorted from cryopreserved PBMC donated by South African adults with new diagnosis of active TB disease (n = 2). (G) Gates were drawn for SGL-CD1b tetramer-positive cells as defined by staining with SGL-CD1b tetramer and negative for mock-loaded CD1b tetramer. To define these gates, we included an SGL-CD1b fluorescence minus one (FMO) control (left), and a T cell line positive control (right), to ensure the gates reliably captured cells that stain with SGL-CD1b tetramer. In this experiment, we included an antistreptavidin antibody conjugated to the same fluorochrome as the SGL-CD1b tetramer to increase the tetramer MFI. (H) SGL-CD1b tetramer positive T cells in the blood were single-cell sorted from cryopreserved PBMC donated by South African adults with new diagnosis of active TB disease (n = 2). This single cell sort was also indexed, meaning that the MFIs for each channel were saved for each single cell, and this information was combined with the TCR assignments to assign a co-receptor group to a cell. The negative population (black) is visualized here using a no tetramer control for each individual (TB-1124, left; TB-1127, right). The sorted cells (red) are visualized within the sort gate (n = 184 per donor).

**Supplemental Figure 2. Differences in functional avidity between SGL-specific T cell lines.** A01 and A05 T cell lines are specific for SGL and examined by multicolor flow cytometry. (A) Tetramer-positive cells within the A05 T cell line express CD4, and lack CD8 and CD8 expression (black). Tetramer-positive cells within the A01 T cell line express CD8 and CD8, but lack CD4 expression (white). (B) Boxplots depict the median and interquartile range of the SGL-CD1b MFI of A01 (white) and A05 (black) from four independent experiments. (Mann-Whitney, p = 0.028, n = 4).

**Supplemental Figure 3. Specificity of staining with SGL-CD1b tetramer.** Jurkat cells were transduced with the germline-encoded mycolyl reactive (GEM) TCR (clone 42) (Gras et al., 2016; Van Rhijn et al., 2013). Jurkat cells transduced with the GEM TCR stain reliably with glucose monomycolate (GMM)-CD1b tetramer and a murine TCR-ß chain constant region (mTCRBC) specific antibody (left). These GMM-specific Jurkat cells do not bind SGL-CD1b or mock-loaded CD1b tetramer (middle, right).

**Supplemental Figure 4. Functional avidity of CD4, CD8, and DN T cells transduced with a GMM-specific TCR.** Flow plot depicts the percent of transduced T cells that are CD4, CD8, or DN. Boxplot depicts the median and interquartile range of the GMM-CD1b or mTCRBC MFI of each CD4, CD8, and DN T cell that was transduced with the GEM TCR. The CD4+ transduced T cells stained with GMM-CD1b tetramer with a 1.29-fold higher MFI than the CD8 GMM-specific T cells, and a 2.00-fold higher MFI than DN transduced T cells (Two-way ANOVA, post-hoc Dunn test, *** = p < 0.0001, n = 107,935). The CD4+ transduced T cells also stained with the anti-murine TCR-ß chain constant region (mTCRBC) antibody with a 1.31-fold higher MFI then CD8+ T cells and a 1.53-fold higher MFI than DN T cells (p < 0.0001 and p < 0.0001, ANOVA with Tukey post-test). The control populations (No Tet and No Ab) were not included in the statistical analysis. Data are representative of two independent rounds of primary T cell transduction with the GEM TCR.

**Supplemental Figure 5. Glycolipid-specific T cells express a diverse TCR repertoire** Variable (V) genes from recovered TCRs ordered by decreasing prevalence in the dataset. For tetramer-sorted cells, V genes were assigned using VDJFasta using IMGT nomenclature. For bulk T cells, V genes were identified using the ImmunoSEQ TCR α assay (Adaptive Biotechnologies). This data set was previously published, and only the two relevant patient samples are analyzed here (DeWitt et al., 2018). Bar plot depicts the percentage of total recovered T cell receptor variable (V) genes identified from (A) SGL-CD1b tetramer-sorted cells from participant TB-1117 (n = 72) and participant TB-1119 (n = 55) (black), compared to the percentage of bulk T cells that utilize that particular V gene (white). (* = 0.01 < p < 0.05, NS = p > 0.05, Fisher’s Exact Test). (B) Bar plot depicts the percentage of total recovered T cell receptor variable (V) genes identified from GMM-CD1b tetramer-sorted cells from participants TB-1117 (n = 30) and TB-1119 (n = 66) (black), compared to the percentage of bulk T cells that utilize that particular V gene (white) (*** = p = 0.0005, Fisher’s Exact Test).

**Supplemental Figure 6. TCR-α chain V gene usage by CD4 and CD8 SGL-specific T cells**. Variable (V) genes from recovered CD4 (black) and CD8 (white) TCRs ordered numerically. For tetramer-sorted cells, V genes were assigned using VDJFasta using IMGT nomenclature. Bar plot depicts the percentage of total recovered T cell receptor variable (V) genes identified from SGL-CD1b tetramer-sorted cells from participants TB-1117 (n = 24), TB-1119 (n = 18), TB-1124 (n = 21), and TB-1127 (n = 19). Co-receptor expression was defined by mRNA in participants TB-1117 and TB-1119. In participants TB-1124 and TB-1127, the tetramer-positive cells were index sorted into the 96-well PCR plate, and CD4 and CD8 expression was defined by staining with anti-CD4 and anti-CD8 antibodies. P-values stated are from Fisher’s exact tests comparing V gene usage in CD4 and CD8 T cells and are unadjusted.

**Supplemental Figure 7. Unsupervised analysis of SGL-CD1b and GMM-CD1b tetramer-sorted cells and tetramer-negative CD3^+^ sorted cells.** (A) Uniform manifold approximation and projection (UMAP) plot of clusters identified using Seurat (Satija et al., 2015). Each dot represents one cell (n = 272 cells). (B) Seven Clusters are uniformly distributed between patients TB-1117 (n = 126) and TB-1119 (n = 149), indicating that all functional clusters are represented by both patient samples and the samples are relatively evenly represented within the dataset. (C) All seven clusters are present among GMM-(n = 96) and SGL-specific T cells (n = 127), suggesting that there may be few functional differences between T cells that recognize both antigens, and that T cells specific for both antigens are evenly represented within the data set (middle). (D) UMAP visualization of all cells colored by CD8 (left) and CD4 (right) expression. Cells that express the respective co-receptor are colored blue and cells that do not express the co-receptor are colored grey. Of note, the cells that express CD4 cluster away from the cells that express CD8. We did not detect CD4 or CD8 transcript in 167 cells. As with most transcriptional profiling methods, failure to detect expression does not necessarily imply lack of protein expression. Thus, cells where we failed to detect co-receptor expression were excluded from the subsequent supervised analysis (n = 105).

**Supplemental Figure 8. Control populations for cytotoxicity assay.** To aid in our cytotoxicity calculation in Figure 5D, we quantified the ratio of K562-CD1b and K562-EV cells in culture. K562-EV and K562-CD1b antigen presenting cells were labeled “low” and “high” with Cell Trace Violet, respectively, and mixed in a 1:1 ratio. These cells were not co-cultured with T cells or lipid antigen. This control enables us to quantify changes in the frequency of K562-CD1b cells that are unrelated to T cell cytotoxic activity to ensure that our “% Cell Death” calculation accurately reflects the reduction of K562-CD1b cells that results from co-culturing with T cells and antigen.

**Supplemental Figure 9. Heatmap of transcription factor expression by sorted T cells.** Heatmap summarizes expression of transcription factors assayed in SGL-CD1b and GMM-CD1b tetramer-positive cells and tetramer-negative T cells (n = 105). Each row represents one cell. Red indicates that the gene was detected, and blue indicates that the gene was not detected. A minimum read count of 5 reads per gene was used to define expression.

