## Supplementary figures and images for "CD4 and CD8 co-receptors modulate functional avidity of CD1b-restricted T cells"

### Supplemental Figure 1

Supplemental Figure 1.

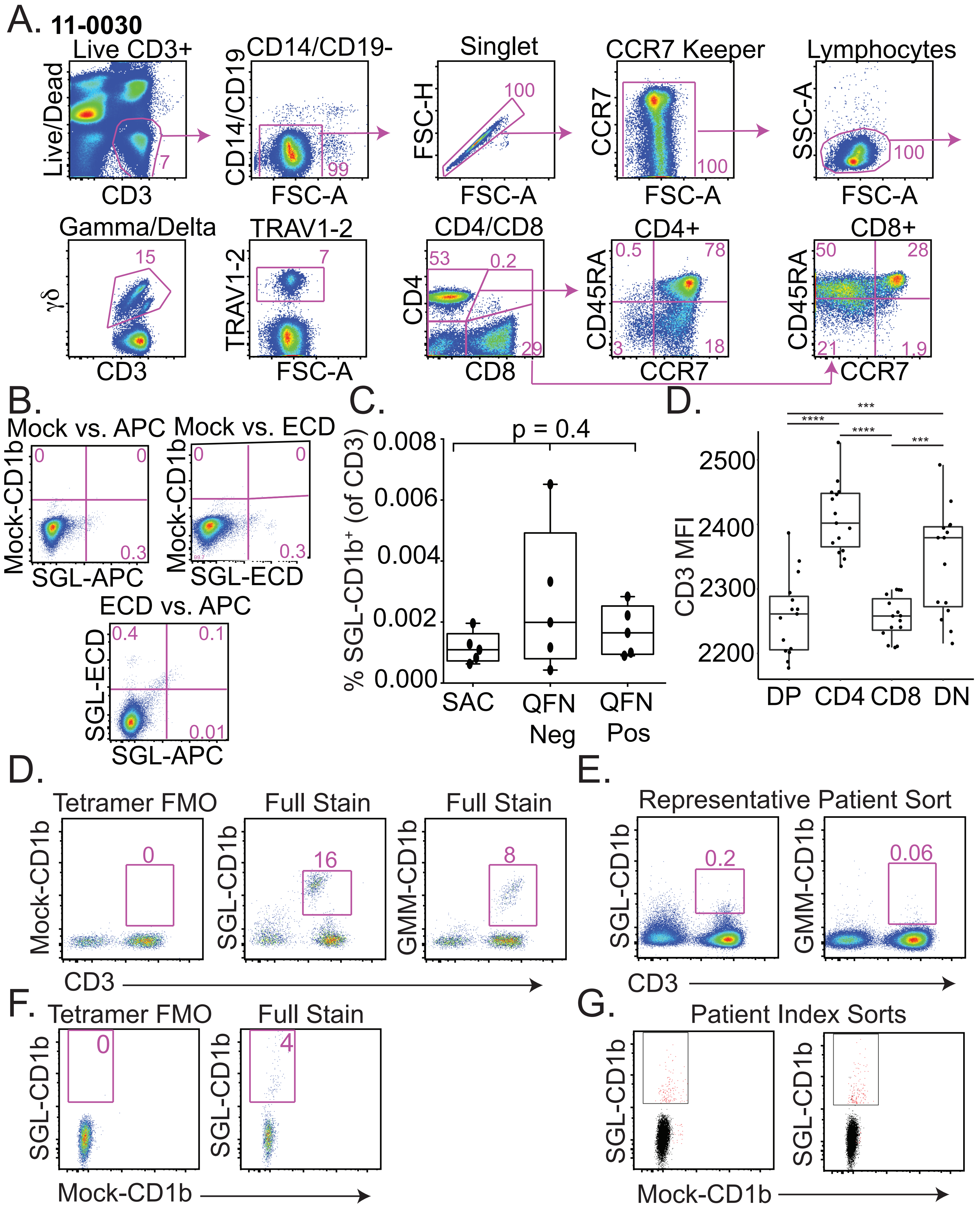

### Supplemental Figure 2

Supplemental Figure 2.

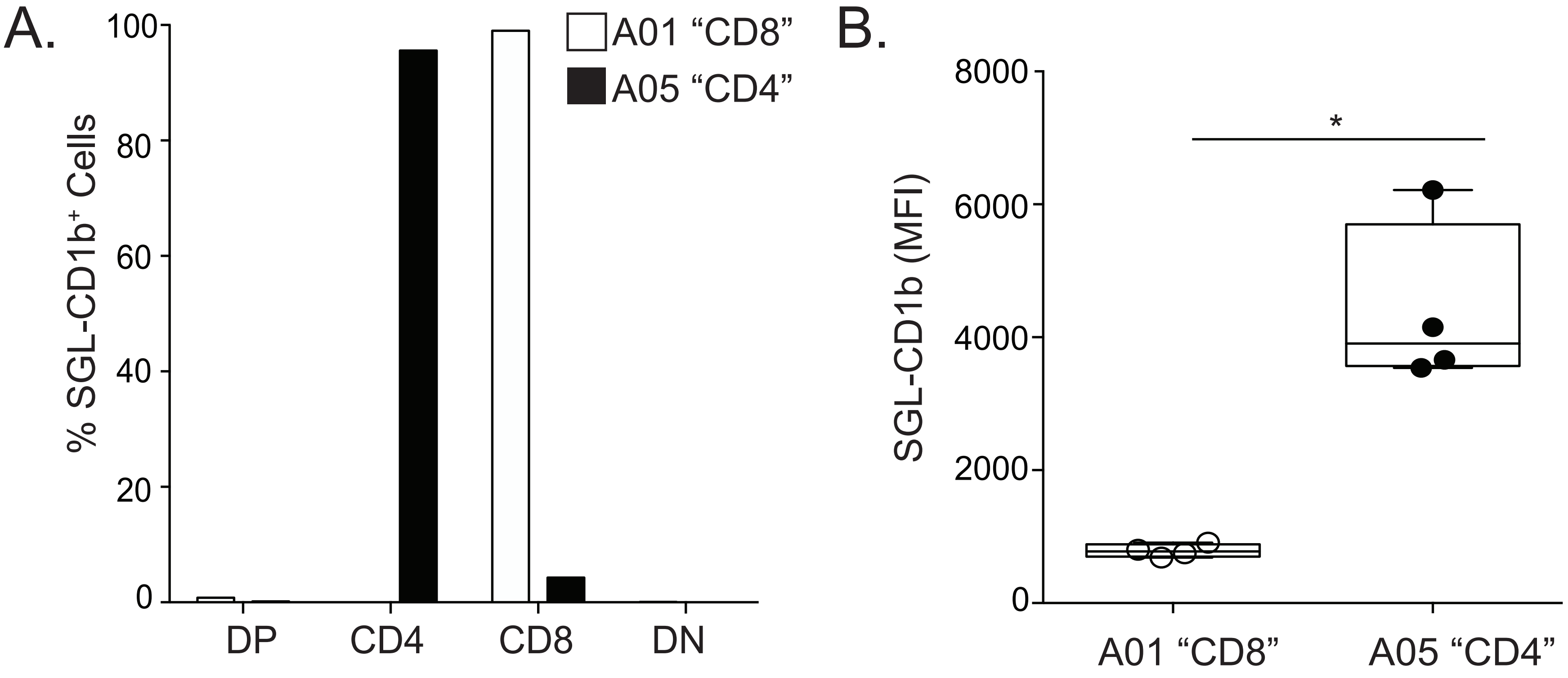

### Supplemental Figure 3

Supplemental Figure 3.

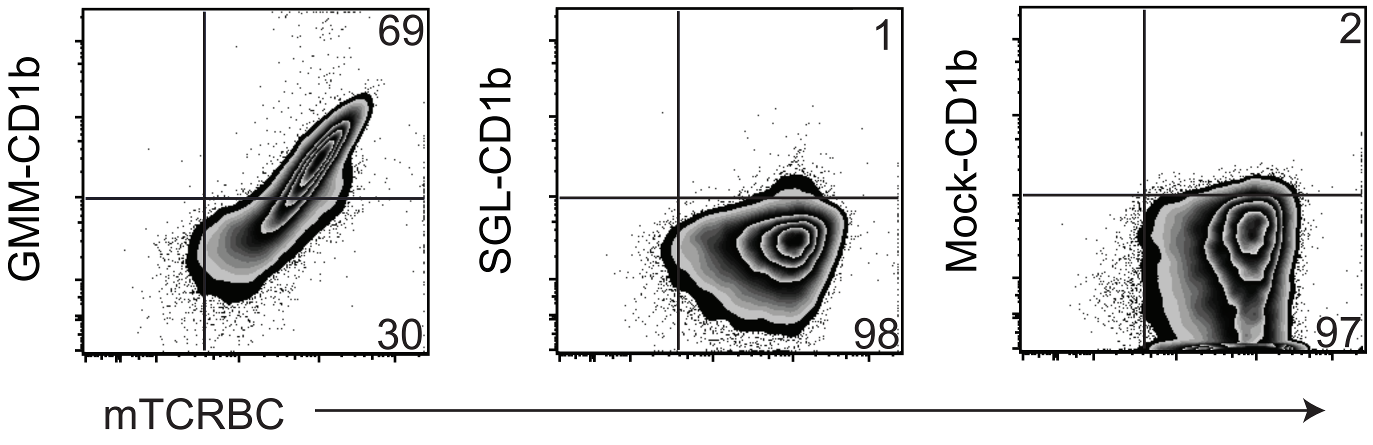

### Supplemental Figure 4

Supplemental Figure 4.

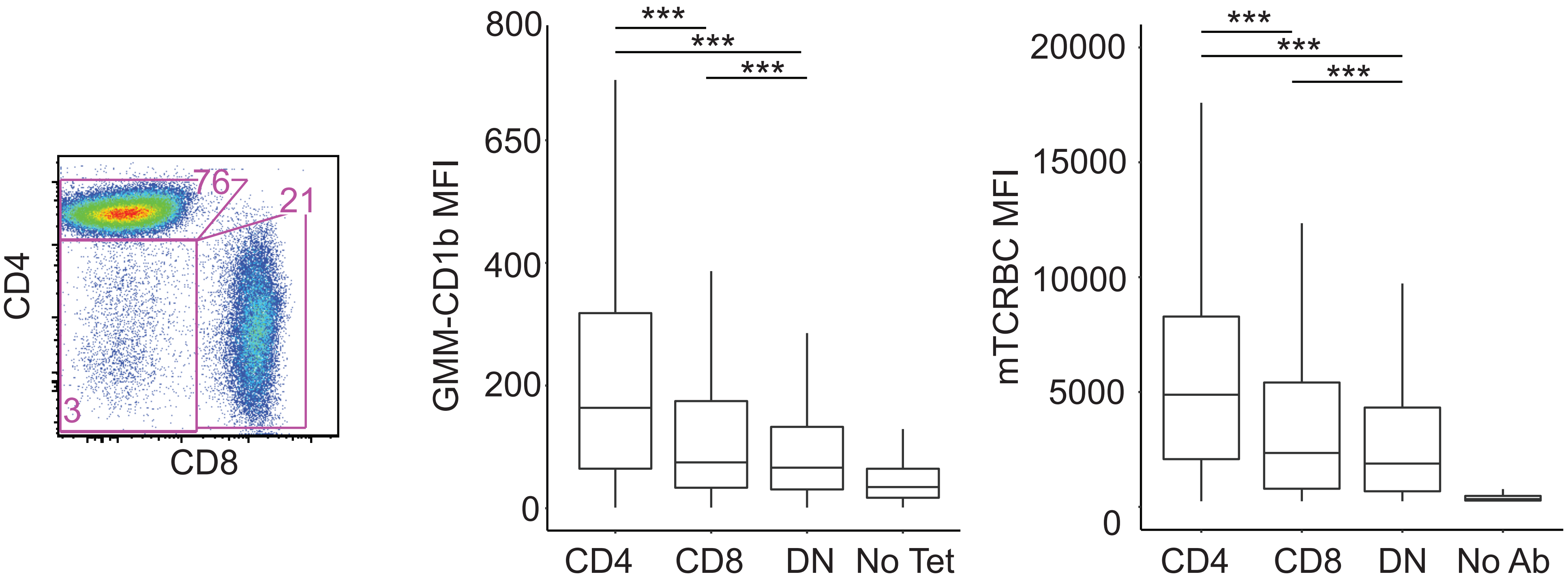

### Supplemental Figure 5

Supplemental Figure 5.

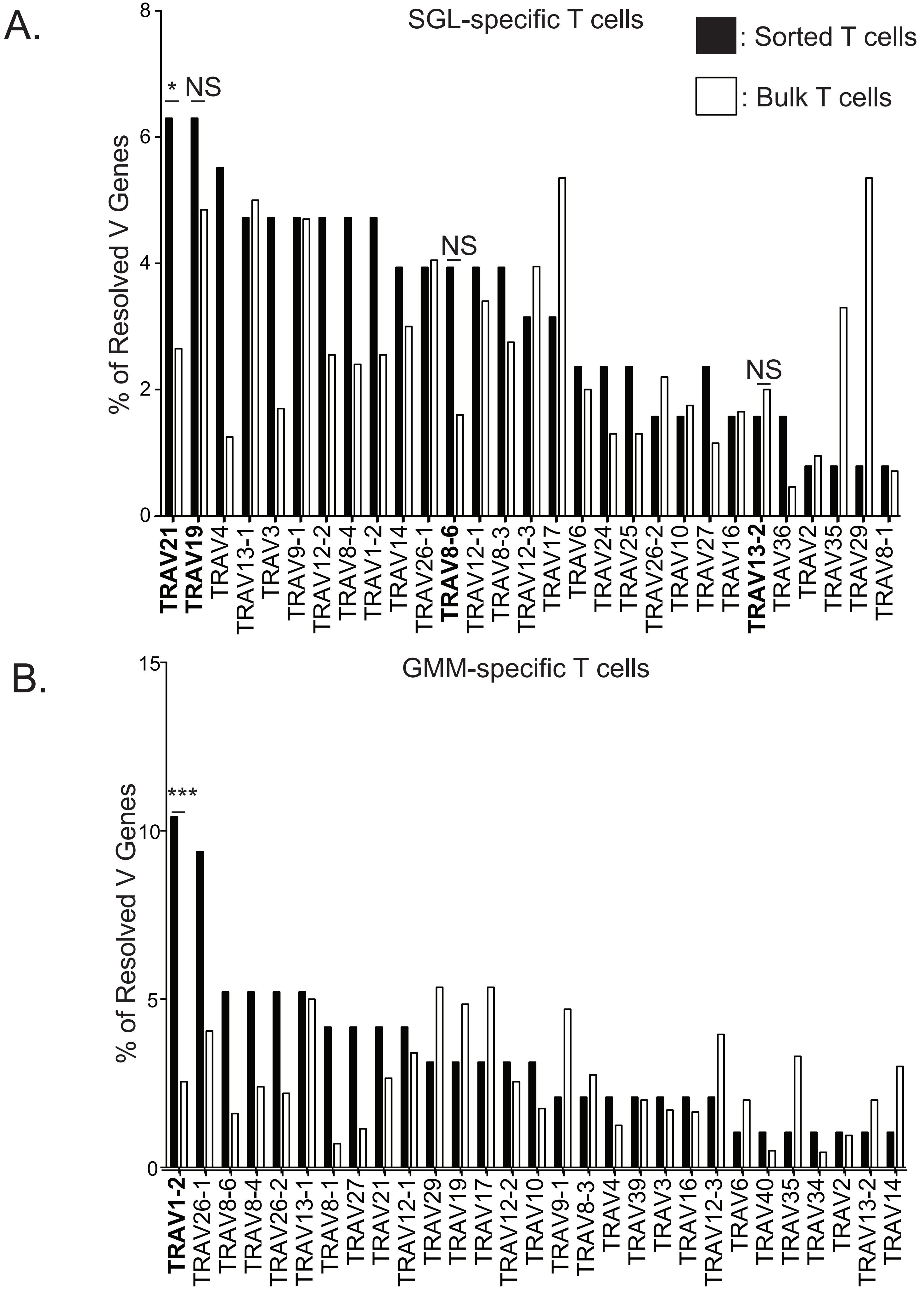

### Supplemental Figure 6

Supplemental Figure 6

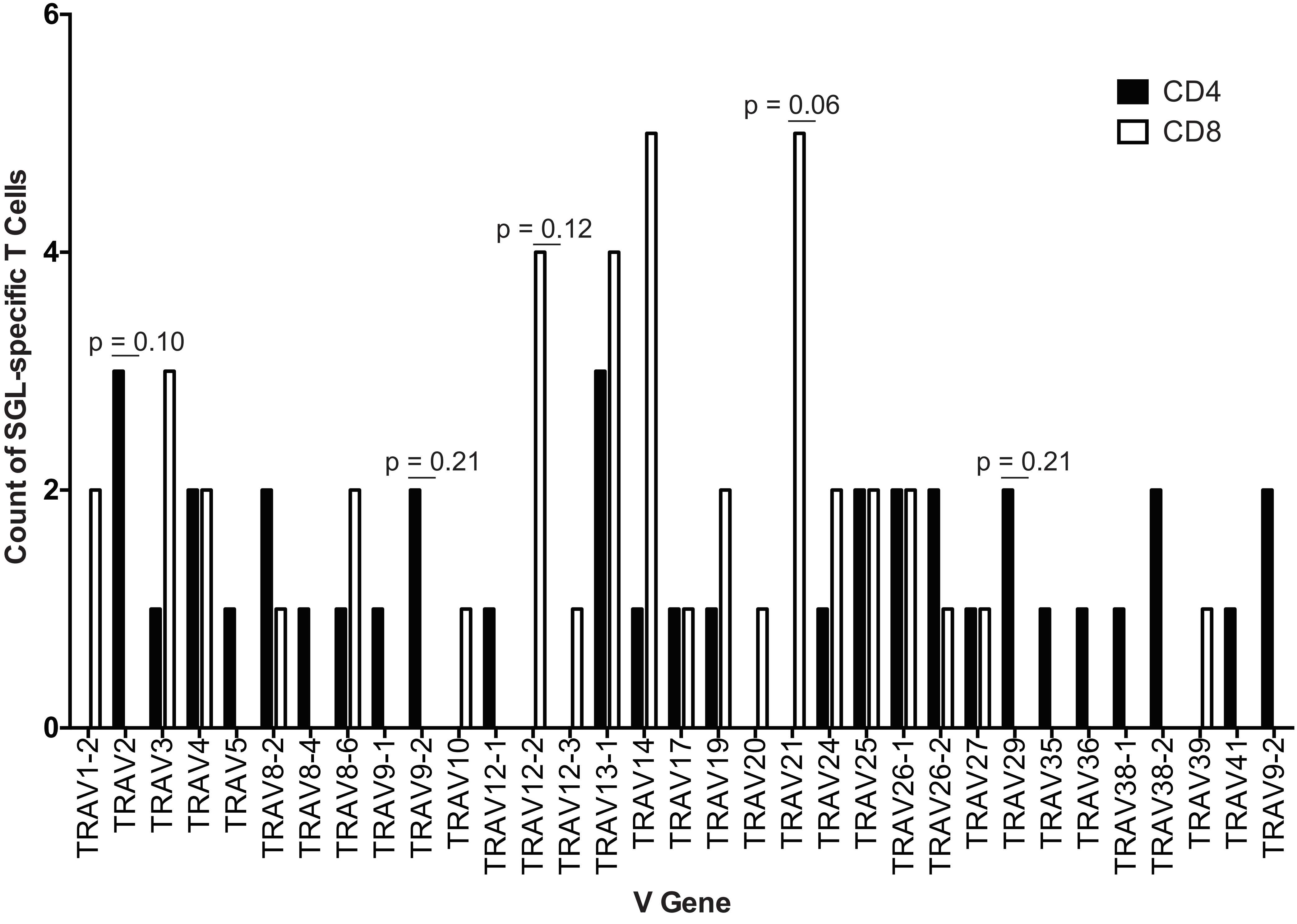

### Supplemental Figure 7

Supplemental Figure 7.

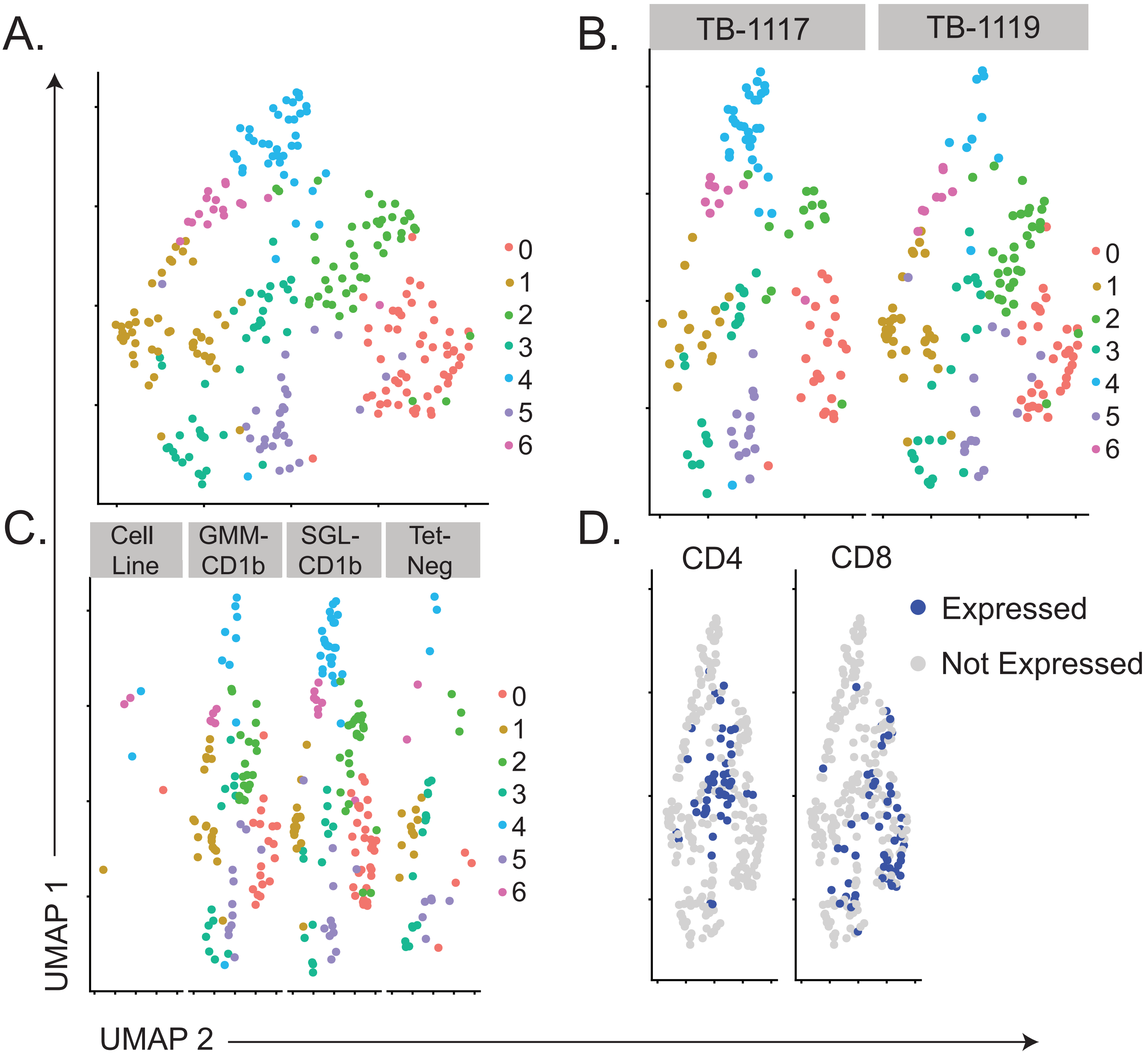

### Supplemental Figure 8

Supplemental Figure 8.

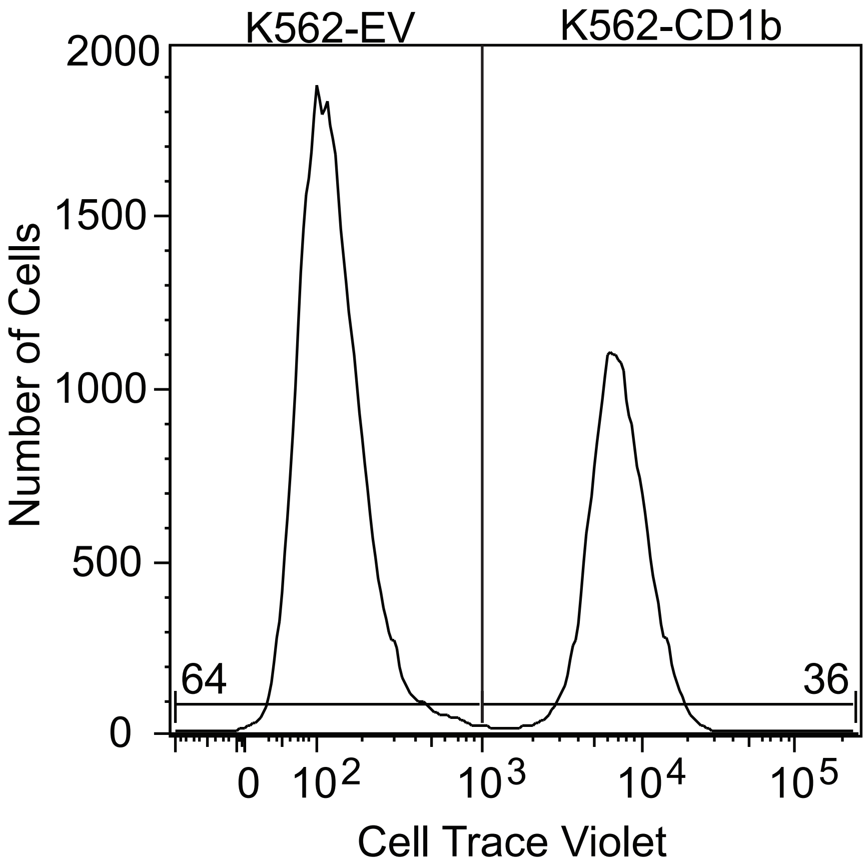

### Supplemental Figure 9

Supplemental Figure 9.

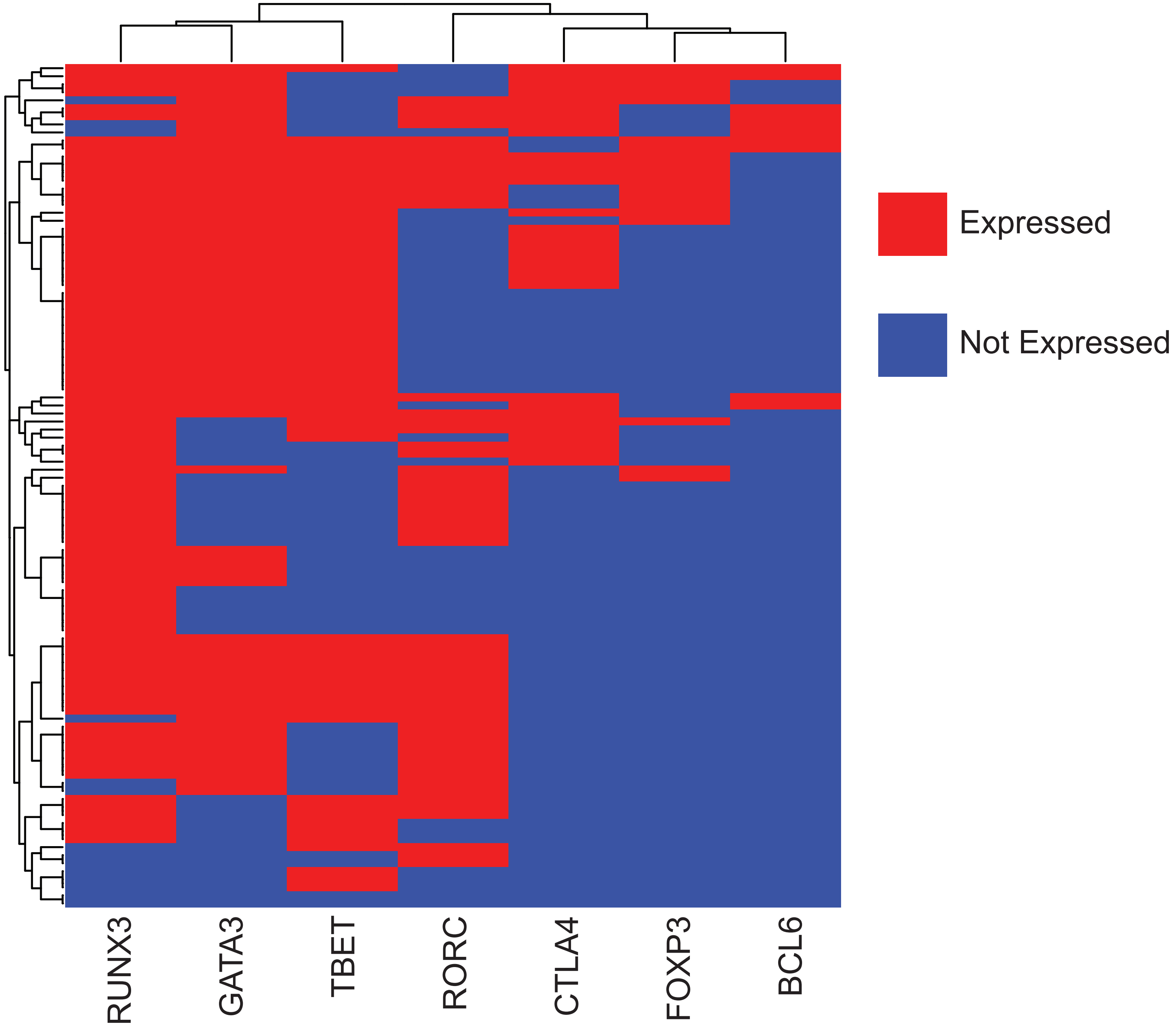
